# Identification of highly immunogenic endogenous dsRNAs from cellular MDA5 filaments

**DOI:** 10.64898/2026.06.07.730679

**Authors:** Jie Chen, ZongChao Mo, Sixing Chen, Xi Wang, Tiffany Hsu, Celia Torres, Kathy Fange Liu, Sun Hur, Qin Li, Roger A Greenberg

**Affiliations:** Department of Cancer Biology, Penn Center for Genome Integrity, Basser Center for BRCA, Perelman School of Medicine, University of Pennsylvania; Philadelphia, PA 19104, USA; Department of Genetics, Penn Center for Genome Integrity, Penn Institute for Immunology & Immune Health (I3H), Perelman School of Medicine, University of Pennsylvania; Philadelphia, PA 19104, USA; Department of Genetics, University of Pennsylvania; Philadelphia, PA 19104, USA; Howard Hughes Medical Institute and Program in Cellular and Molecular Medicine, Boston Children’s Hospital; Department of Biological Chemistry and Molecular Pharmacology, Harvard Medical School; Boston, MA 02115, USA; Program in Cellular and Molecular Medicine, Boston Children’s Hospital; Boston, MA 02115, USA; Department of Biochemistry and Biophysics, Penn Center for Genome Integrity, Penn Institute for RNA Innovation, Perelman School of Medicine, University of Pennsylvania; Philadelphia, PA 19104, USA

## Abstract

ADAR1 converts adenosine to inosine in endogenous double-stranded RNAs (dsRNAs) to prevent excessive MDA5-driven interferon-stimulated gene expression. The source of endogenous immunogenic dsRNAs remains enigmatic because only a small fraction of ADAR1 substrates activate MDA5, and cellular MDA5 filaments have not been isolated. Here, we couple affinity purification of cellular MDA5 filaments with RNA sequencing to define immunogenic endogenous dsRNAs. Greater than 84% of dsRNAs suppressed by combined DDX3X RNA helicase and ADAR1 base-editing activities were present in MDA5 filaments, compared to less than 1% of dsRNA substrates acted on by ADAR1 alone. Dual substrate dsRNAs consisted of inverted repeats embedded in 3’-UTRs with high base-pair complementarity and longer intervening sequences between repeats, with a minor contribution coming from intermolecular dsRNAs formed by sense and antisense transcripts. Moreover, the majority of dual substrate immunogenic dsRNAs were hyperedited in DDX3X mutant cancers. This reveals the identity of endogenous immunogenic dsRNAs and quality control mechanisms underlying their suppression.

## Main

Melanoma differentiation-associated gene 5 (MDA5) and retinoic acid-inducible gene I (RIG-I) form filamentous structures on their cognate dsRNA ligands^1–5^. Active filament formation promotes aggregation of the MAVS protein on the mitochondrial surface and signaling through IRF3 and NF-κB transcription factors to elicit interferon-stimulated gene (ISG) expression in response to either foreign or endogenous immunogenic dsRNAs^6^. MDA5-substrate availability is negatively regulated by ADAR1, which converts adenosine to inosine on dsRNA to prevent MDA5 activation^7–9^. Loss of function mutations to ADAR1 result in the pediatric autoimmune syndrome Aicardi-Goutières, characterized by high interferon expression, brain calcifications and features of lupus^10^. Activating mutations in MDA5 that reduce its sensitivity to dsRNA mismatches or A-to-I base changes produce a similar syndrome^11,12^. Aicardi-Goutières in both settings occurs in the absence of known viral or bacterial infections, indicative of an endogenous source of sterile inflammation. In agreement, enzymatically inactive ADAR1 mutant mice exhibit embryonic lethality that is suppressed by MDA5 deficiency^7,9,13,14^. Complete deletion of ADAR1 p150 isoform produces a more severe lethality that is reversed by combined loss of both MDA5 and protein kinase R (PKR)^15,16^. This increased severity is thought to arise because, in addition to RNA editing, ADAR1 sequesters dsRNA from other sensors, such as PKR^17,18^. Collectively, these observations indicate that lethal inflammation from basal production of endogenous immunogenic dsRNA ligands is prevented by constitutively active quality control mechanisms.

Additional regulatory mechanisms for MDA5 engagement with dsRNAs have been suggested. Deficiencies in DDX3X and DHX9 RNA helicases were reported to enhance ISG responses. Combined DHX9 and ADAR1 loss enhanced dsRNA accumulation and ISG responses^19,20^. DDX3X RNA helicase knockdown also enhanced ISG responses, and DDX3X loss was reported to reduce ADAR1 editing activity in cells^21^. However, the immunogenic MDA5 ligands that arise in cells lacking RNA helicase and ADAR1 base editing activities have not been identified.

The identity of the endogenous dsRNA species that activate MDA5 has remained elusive for technical reasons. MDA5 exists as a monomeric species with relatively high non-specific affinity for a variety of nucleic acids, while the active form is a filament in complex with long dsRNAs^1,22^. RNA sequencing of ADAR1 A-to-I editing represents an indirect measure of the cytoplasmic dsRNA species and has been used to infer the nature of activating MDA5 ligands^23^. However, this type of indirect analysis cannot pinpoint the active MDA5 ligands because ADAR1 knockout experiments indicate that less than 2% of all edited dsRNAs are responsible for MDA5 activation^13,24–26^. This is likely because the MDA5 protein is sensitive to mismatches in dsRNA as well as A-to-I base substitutions^27,28^, thus limiting its activation by endogenous dsRNAs. Moreover, methods to purify endogenous MDA5 filaments from cells have not been reported, making direct identification of active dsRNA ligands a technically intractable issue.

Here we isolate the active MDA5 filament, enabling identification of its endogenous dsRNA ligands. Coupled with CRISPR screening approaches and analyses of adenosine to inosine editing profiles, we determine that DDX3X RNA helicase and ADAR1 editing activities cooperate to suppress highly active MDA5 ligands. Greater than 90% of these dual-substrate dsRNAs were inverted repeat Alu elements embedded within the 3’-UTR (untranslated region) of RNA polymerase II transcripts that contain high repeat complementarity, longer distances between inverted repeat elements, and lower predicted free energy. These findings reveal the identities of endogenous immunogenic dsRNAs and the basis for their suppression by biochemically distinct quality control mechanisms.

## Results

### DDX3X RNA helicase activity limits irradiation (IR)-dependent inflammatory signaling

We performed a targeted array-based CRISPR Cas9 screen to identify proteins involved in the regulation of DNA-damage-induced dsRNA pattern recognition mediated ISG induction. The screen was conducted in hTERT RPE-1 p53 KO cells, which express MDA5 and RIG-I but are cGAS deficient (Fig.1a,b). Prior reports revealed that ATR inhibition (ATRi) in p53 null cells could activate a strong ISG response that relied on dsRNA sensors RIG-I and MDA5 but was independent of cGAS^29–31^. ISG markers RIG-I, ISG56, and IFITM1 were significantly increased at 3 days following irradiation and ATR inhibition (IR, 10 Gy + ATRi). Guide RNAs (gRNAs) targeting inflammatory signaling mediators IRF3, IKBKG, RELA, or IFNAR1, abolished ISG upregulation, whereas gRNAs targeting MDA5 or RIG-I reduced ISG induction by approximately 50%. Guide RNAs targeting STING had no effect on ISG expression, in agreement with cGAS not being expressed in RPE-1 cells.

**Fig. 1.**
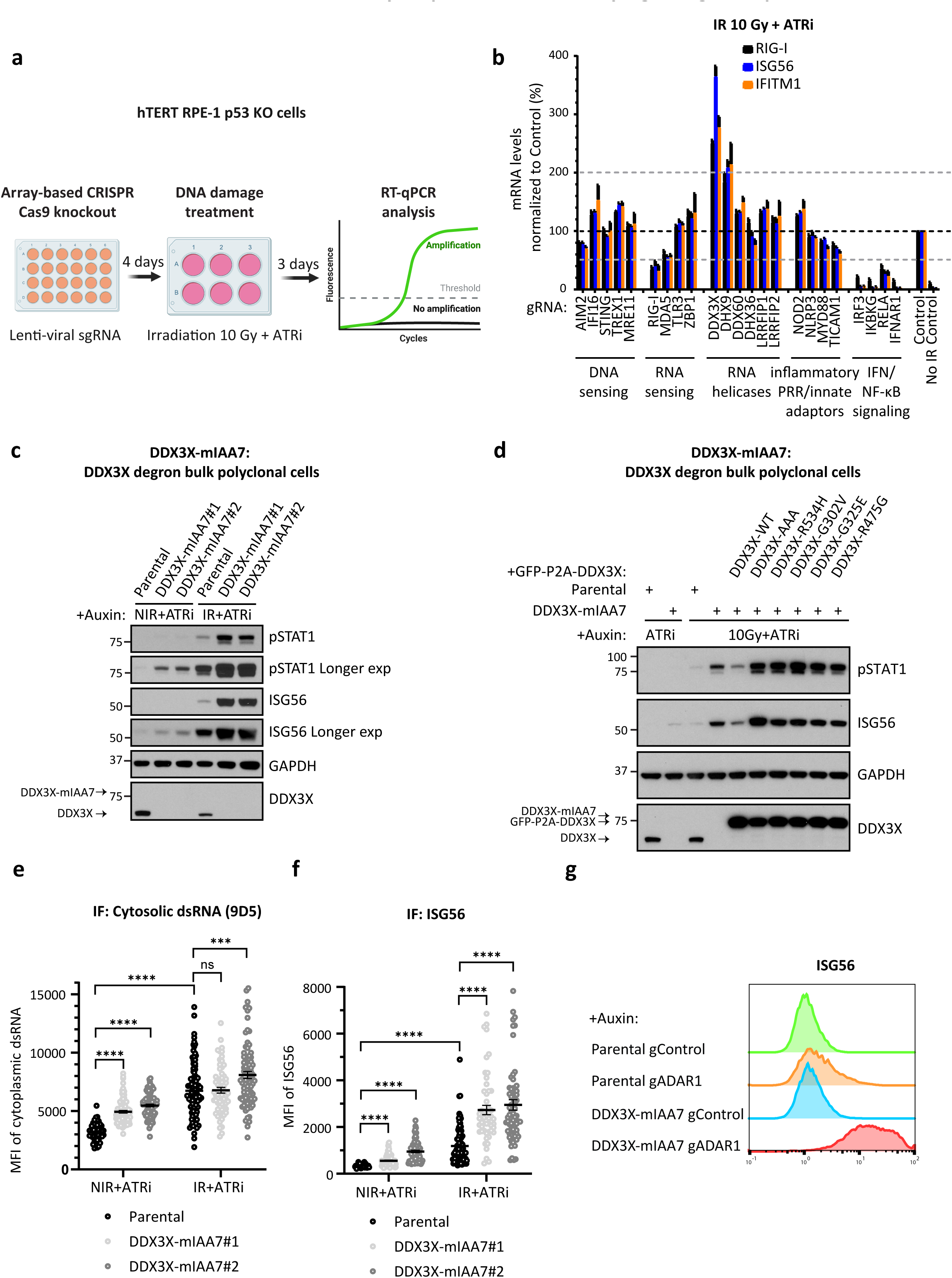
Loss of DDX3X RNA helicase activity amplifies inflammatory signaling in response to DNA damage or ADAR1 deficiency. **a**, Schematic showing CRISPR-Cas9 screening workflow. Created with BioRender. **b**, Cells were treated as depicted in **a** and analyzed by RT-qPCR analysis. **c**, Cells were treated with 1 mM auxin in the presence of 2.5 µM ATRi for 3 hours, followed by 10 Gy irradiation (IR) or no irradiation (NIR). Cells were cultured for another 3 days in the presence of 1 mM auxin and 2.5 µM ATRi before western blot analysis. **d**, Cells were treated with 2 µg/ml doxycycline (DOX) for 24 hours, then incubated with 2 µg/ml DOX, 1 mM auxin and 2.5 µM ATRi for 3 hours before 10 Gy irradiation (IR) or no irradiation (NIR). Cells were cultured for another 2 days in the presence of 2 µg/ml DOX, 1 mM auxin and 2.5 µM ATRi before western blot analysis. **e**, **f**, Mean Fluorescence Intensity (MFI) of cytosolic dsRNA detected by the 9D5 antibody or anti-ISG56 in cells following auxin induced DDX3X depletion, 10 Gy IR + ATRi or combined DDX3X deficiency and IR + ATRi. **g**, Four days post lentiviral guide RNA infection, cells were treated with 1 mM auxin for 2 days before collection for flow cytometry analysis. Flow cytometry for ISG56 levels following DDX3X degradation, CRISPR deletion of ADAR1 (gADAR1), or combined DDX3X and ADAR1 deficiency reveals greater than additive increases in ISG expression following loss of both DDX3X and ADAR1.

The screen identified RNA helicases DDX3X and DHX9 as suppressors of ISG expression (Fig. 1b and Extended Data Fig.1a), with loss of DDX3X producing the greatest increases in all three ISGs monitored. Since DDX3X is an essential gene^32,33^, we generated a DDX3X-mIAA7 degron cell line^34,35^ to enable its inducible depletion. Both the DDX3X-mIAA7 degron knock-in efficiency (>95%) and auxin-induced depletion efficiency were very high (Extended Data Fig.1b-d), allowing us to perform experiments in bulk polyclonal cell populations. Tagging DDX3X with mIAA7 using two different gRNAs neither affected its basal expression levels nor impacted DNA damage-induced inflammatory signaling in the absence of auxin (Extended Data Fig.1c-e). However, upon auxin-induced DDX3X depletion, we observed a consistent increase in inflammatory signaling in both IR+ATRi treated and ATRi-only-treated cells (Fig.1c and Extended Data Fig.1f). Phosphorylated IRF3 and p65 were mildly elevated, indicating increased activation of both signaling pathways (Extended Data Fig.1f). Given that DDX3X is an RNA helicase^36,37^, we tested whether the unwinding activity was required for its function. Wild-type DDX3X, but not unwinding-deficient mutants^38–40^, suppressed the elevated inflammatory signaling induced by DDX3X depletion, highlighting the importance of its helicase activity in limiting inflammatory signals (Fig.1d). DDX3X mutants increased ISG expression beyond that observed from DDX3X degradation alone. This is likely because auxin induced degradation of DDX3X is incomplete (Extended Data Fig.1d) and overexpression of an inactive mutant further suppresses the activity of the residual DDX3X-mIAA7 protein.

These findings are consistent with a model that DDX3X depletion cooperates with DNA damage to increase immunogenic dsRNA production. To test this hypothesis, we monitored cytoplasmic dsRNA levels in DDX3X deficient cells at baseline and after IR. We employed saponin permeabilization^41^ and used the 9D5 anti-dsRNA antibody, which is more sensitive than the J2 anti-dsRNA antibody in detecting cytoplasmic dsRNAs^42^ (Extended Data Fig.2a). DDX3X depletion alone increased cytosolic dsRNA levels as did IR, consistent with independent contributions from each perturbation (Fig.1e, and Extended Data Fig.2b,e,f), and that DDX3X actively suppresses dsRNA formation independent of DNA damage. In accordance, DDX3X degradation combined with IR elevated ISG induction in ATRi + IR treated cells in a greater than additive manner (Fig.1f, and Extended Data Fig.2g,h). We further hypothesized that DDX3X would also play an important role in other conditions known to accumulate dsRNA^43^. In agreement, individual depletion of either DDX3X or ADAR1 led to a mild increase in ISG56 expression, while simultaneous depletion of DDX3X and ADAR1 caused a synergistic elevation in ISG56 expression (Fig.1g and Extended Data Fig.2c,d). ISG induction in ADAR1- and DDX3X-deficient cells was largely mediated by the dsRNA sensor MDA5 (Extended Data Fig.2d), indicating that DDX3X and ADAR1 cooperate to prevent immunogenic dsRNA MDA5 ligands.

### DDX3X depletion induces the formation of specific dsRNAs

ADAR1 editing is an established means of suppressing the formation of MDA5 activating dsRNA ligands, and DDX3X has been proposed to stimulate its editing activity^21^. Given that DDX3X helicase activity represents a distinct biochemical mechanism of limiting dsRNA formation, we instead postulated that DDX3X would function to limit the pool of ADAR1 substrates. This model predicts that ADAR1 would buffer the loss of DDX3X to limit MDA5 activating dsRNA ligand accumulation. We investigated this function at higher resolution by performing RNA editing analysis on purified RNA from fractionated cytoplasmic lysates^44^. Inosine is structurally similar to guanosine and is detected by A-to-G changes on RNA sequencing reads. ADAR1 editing produces A-to-I substitutions exclusively on dsRNA substrates, making recurrent A-to-G sequencing read alterations as proxies for dsRNA formation. Analyzing ADAR1 editing clusters of greater than five edits within close proximity to each other significantly improves signal-to-noise ratio for detecting long, potentially immunogenic dsRNAs^24^ (Fig.2a). Consistent with the model that DDX3X suppresses the formation of ADAR1 substrates, we observed approximately 1.6-fold increases in the total amount of highly edited dsRNAs in DDX3X deficient cells (Fig.2b). The increase in edited dsRNAs following DDX3X depletion was not due to changes in the level of RNA harboring dsRNA, as there was no overall significant difference in dsRNA expression levels between control and DDX3X depleted cells (Fig.2c). The subtle difference in ADAR1 expression level between control (10.5 TPM) and DDX3X depleted (12.1 TPM) cells was also not sufficient to explain the increased editing. Instead, DDX3X depleted cells contained more than twice as many edited dsRNA species than the control parental cells (Fig.2d), with an increased prevalence of dsRNA formation mostly found in 3’-UTR regions of mRNAs (Extended Data Fig.3a). More than 90% of edited dsRNAs occurred on SINE/Alu elements in the parental and DDX3X degraded cells in agreement with reported editing profiles^45–47^, with some additional dsRNAs observed on other transposable repeat sequences (Extended Data Fig.3b,c). This raises the possibility that DDX3X suppresses a specific class of highly immunogenic Alu elements.

**Fig. 2.**
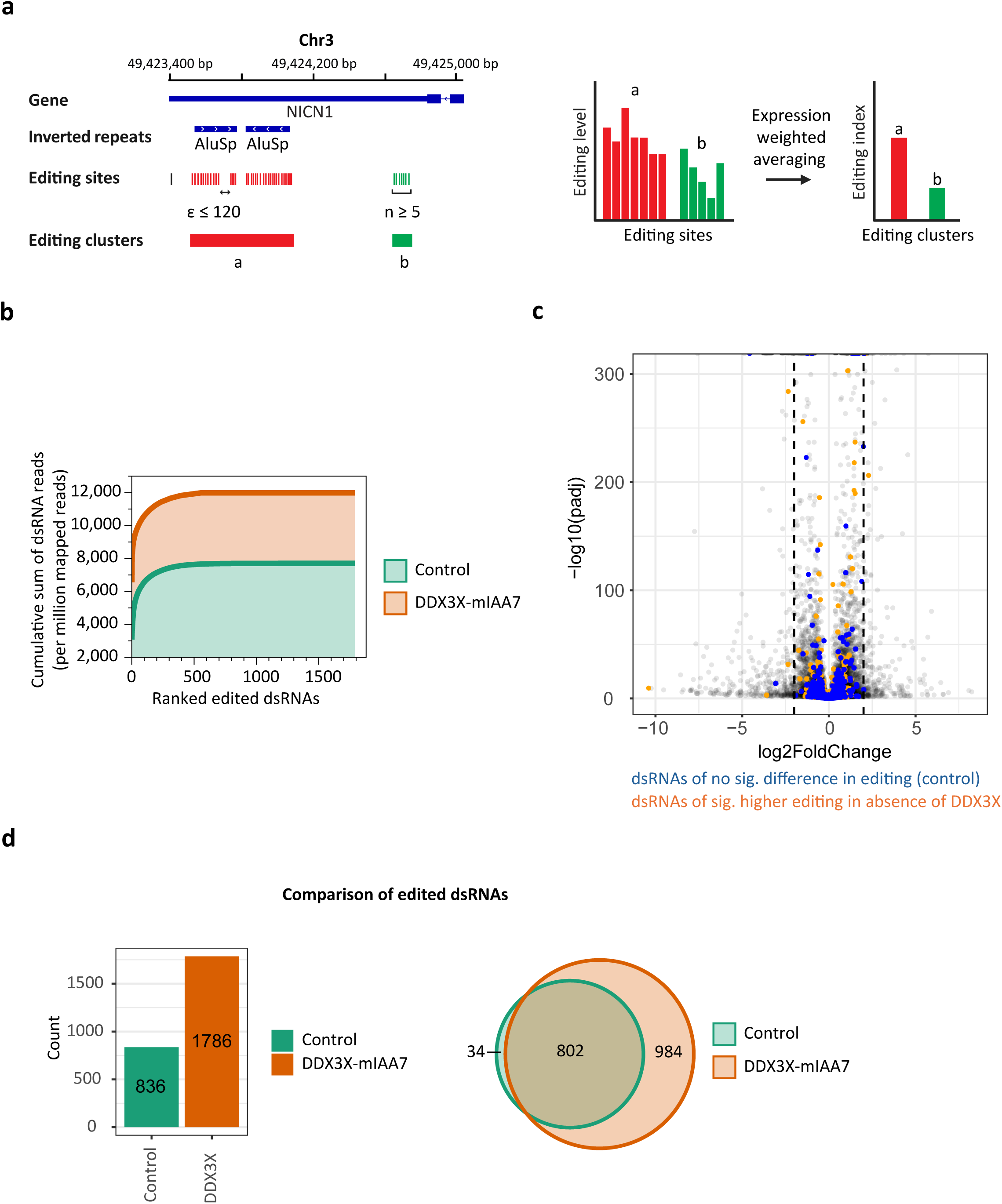
DDX3X depletion increases the formation of dsRNA substrates for ADAR1 editing. **a**, Left panel: Example of editing clusters. Proximal editing sites were grouped as one cluster if (1) the number of editing sites is at least 5 and (2) the distance between any two adjacent sites is less than 120 nt (see **Methods** for details). Right panel: Schematic diagram of converting editing levels measured at individual sites of a cluster to the editing index of an entire editing cluster (see **Methods** for details). Created with BioRender. **b–d**, Cells were treated with 1 mM auxin for 2 days to induce DDX3X degradation (DDX3X-mIAA7), followed by cytoplasmic RNA extraction and strand-specific RNA-seq. **b**, Comparison of dsRNA abundance between DDX3X depleted (DDX3X-mIAA7) and control cells. We define dsRNAs using editing clusters and the abundance was measured as edited reads normalized by per million mapped reads. **c**, Differential expression analysis between DDX3X depleted and control cells. Colored dots represent dsRNAs either differentially edited upon DDX3X depletion (orange, also see Fig. 4a) or non-differentially edited (blue). **d**, Comparison of dsRNA loci identified by *de novo* RNA editing analysis (see **Methods** for details). Only hyper-edited dsRNAs not included in the RADAR editing database were used for comparison.

### Identification of immunogenic dsRNA ligands activating MDA5

Structural analysis of RIG-I-like receptors (RLRs) revealed that MDA5 and RIG-I filaments are stabilized by cognate E3 ubiquitin ligases that specifically act on the filamentous form of each RLR through avidity interactions with oligomeric MDA5 or RIG-I. The E3 ligase TRIM65 specifically recognizes MDA5 filaments through bivalent interactions but not inactive MDA5 monomers^48^. This finding provides the rationale for using recombinant TRIM65 to specifically purify oligomeric MDA5 filaments and sequence its associated dsRNAs (Fig.3a and Extended Data Fig.4a). Purified recombinant Streptavidin-Binding Peptide (SBP)-tagged TRIM65 fragments inclusive of the coiled-coil and PRY-SPRY domains were used to pull down active MDA5 filaments in DDX3X deficient cells with concurrent ADAR1 deletion or treatment with IR + ATRi. Purifications in MDA5 KO cells served as a control for nonspecific RNA binding.

**Fig. 3.**
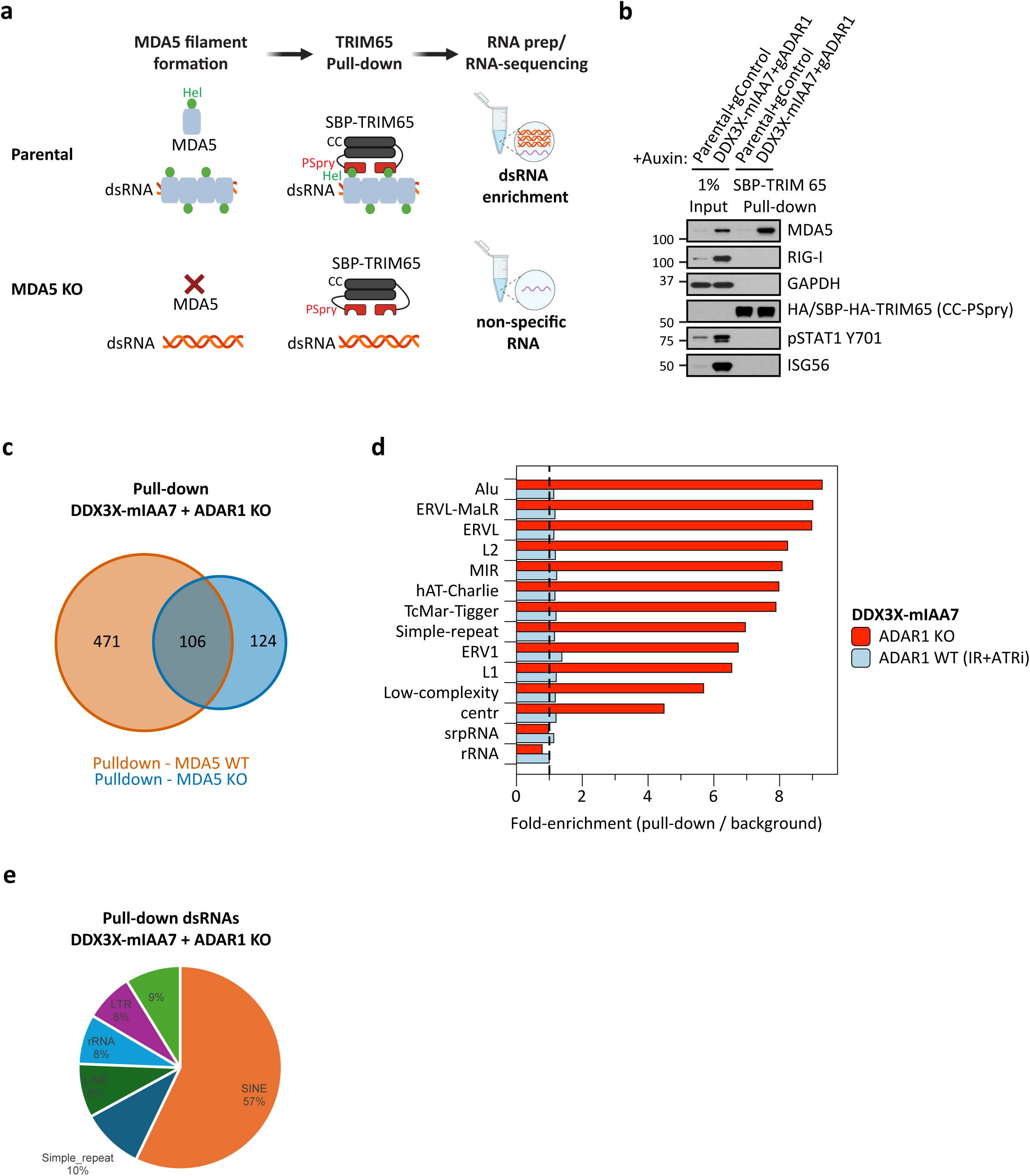
Discovery of MDA5 associated dsRNA ligands in cells lacking DDX3X and ADAR1. **a**, Schematic of the SBP-TRIM65 (streptavidin binding peptide tagged TRIM65) pull-down strategy used to identify endogenous immunogenic dsRNA ligands associated with MDA5 filament formation, comparing parental and MDA5 knockout cells. dsRNAs are selectively enriched in the presence of MDA5. Created with BioRender. **b-e**, Cells were infected with lentiviral guide RNA targeting indicated genes. 4 days post infection, cells were treated with 1 mM auxin for 2 days before cytoplasmic fraction for TRIM65 pull-down. **b**, Western blot analysis of input and TRIM65 pull-down samples showing activation of downstream interferon signaling markers and MDA5 association with TRIM65. **c**, Venn diagram showing overlap of RNA peaks recovered from TRIM65 pull-downs in MDA5 wild-type versus MDA5 knockout cells in a DDX3X degradation (DDX3X-mIAA7) and ADAR1 knockout (ADAR1 KO) background. **d**, Fold enrichment of repetitive elements in TRIM65 pull-downs from the indicated conditions (DDX3X depleted, ADAR1 KO, or DDX3X depleted, ADAR1 WT, IR+ATRi), normalized to the corresponding MDA5 KO background pull-down and grouped by repeat class. **e**, Relative contribution of repetitive elements classes to dsRNAs enriched in TRIM65 pull-downs.

Recombinant TRIM65 demonstrated specific interaction in cellular lysates with MDA5 but not with RIG-I or any other proteins tested (Fig.3b and Extended Data Fig.4b,c). Approximately 2% of the total MDA5 was precipitated by TRIM65 under these conditions. The interaction between TRIM65 and MDA5 remained intact even after high-concentration RNase A digestion (1 mg/ml), which was used to reduce the presence of associated nonspecific RNAs and further enhance the signal-to-noise ratio (Extended Data Fig.4b,c). In DDX3X and ADAR1 co-depleted cells, approximately 18% of RNAs purified were present in the MDA5 KO control. These overlapping RNAs were excluded from further analysis (Fig.3c). We observed a 4.5- to 9.5-fold enrichment of different classes of inverted repeat transposons and endogenous retroviral dsRNAs following ADAR1 and DDX3X co-depletion (Fig.3d). 57% of the TRIM65 purified RNAs were SINE/Alu elements upon ADAR1 and DDX3X co-depletion (Fig.3e). IR + ATRi did not show similar enrichments of Alu and endogenous retroviral elements (Extended Data Fig.4d), suggesting that DNA damage cooperates with DDX3X deficiency to produce different immunogenic dsRNA species^49^. Notably, 43% of the total TRIM65 purified RNAs following exclusion of the MDA5 KO background were derived from other RNA classes and did not display A-to-I editing. The non-edited MDA5 ligands likely arise from ADAR1 deletion, which is known to cause a redistribution of dsRNAs to other sensors^17,18^.

### DDX3X helicase and ADAR1 editing activities suppress the formation of highly active MDA5 ligands

We reasoned that dsRNAs that are differentially edited by ADAR1 in DDX3X deficient cells would represent a group of highly active MDA5 ligands. To identify these dual-substrate dsRNAs, we performed an unbiased analysis of RNA editing index and identified 292 dsRNAs that were significantly more edited by ADAR1 upon loss of DDX3X (hereafter referred to as *differentially edited dsRNAs*, DEDs; Fig.4a,e). TRIM65-mediated purification of active MDA5 filaments from ADAR1 and DDX3X co-depleted cells recovered 246 of the 292 DEDs, corresponding to ∼84% of all DEDs identified by editing analysis (Fig.4a,b,e,f). To assess whether DEDs are preferentially engaged by MDA5, we examined MDA5 occupancy profiles across DED and non-differentially edited (non-DED) dsRNAs. DEDs exhibited strong and reproducible enrichment in TRIM65-mediated MDA5 pulldowns across their length in wild-type cells, whereas this enrichment was abolished in MDA5 knockout control cells (Fig.4c). In contrast, non-differentially edited dsRNAs showed minimal MDA5 enrichment and no discernible occupancy pattern, despite comparable detection in input samples (Fig.4d). These results indicate that DEDs are selectively incorporated into active MDA5 filaments, rather than representing a general consequence of dsRNA abundance or editing *per se*.

**Fig. 4.**
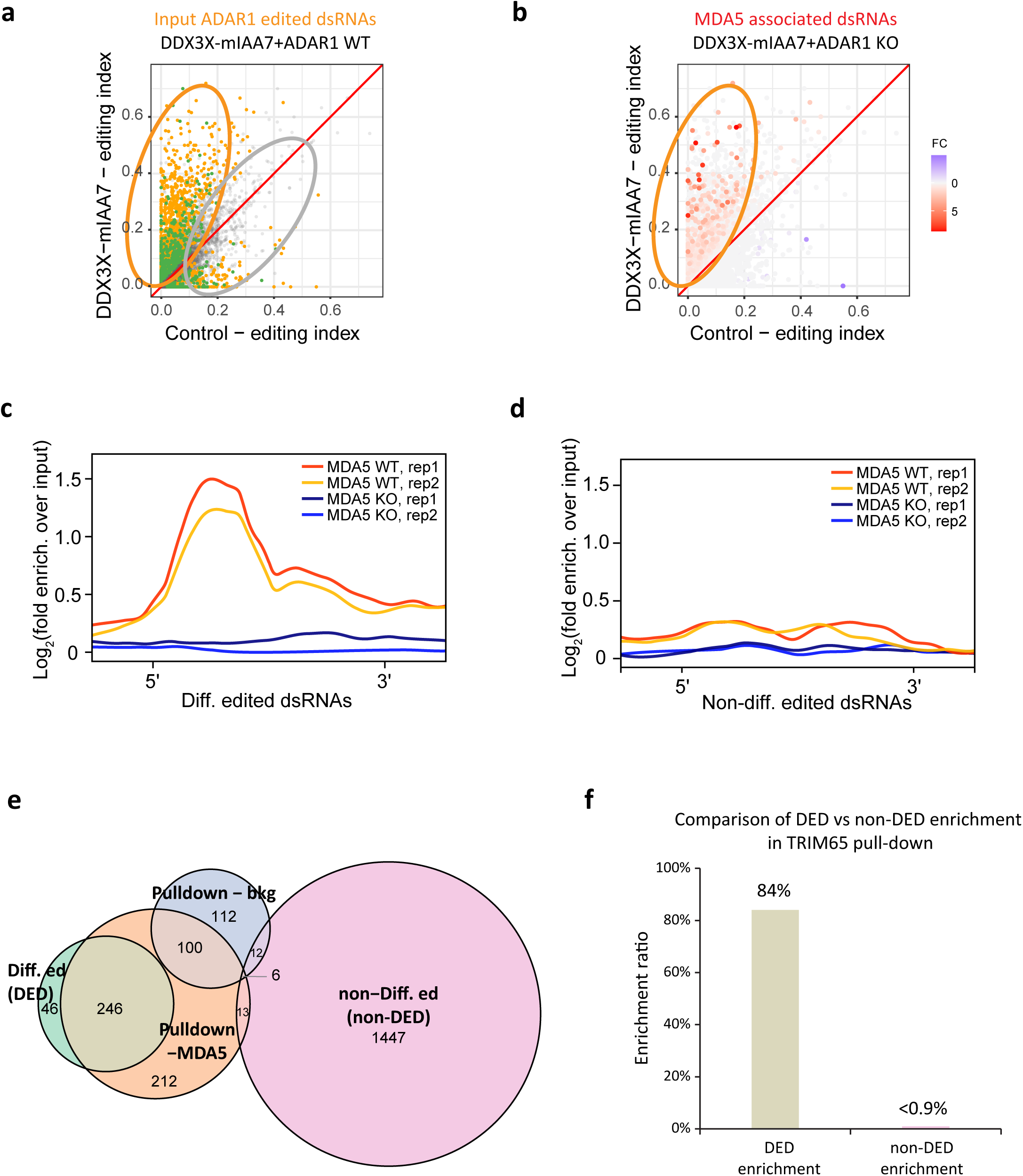
Differentially edited DDX3X dsRNA substrates are highly enriched in MDA5 filaments. **a**, Comparison of dsRNAs editing identifies DDX3X-dependent dsRNAs. Editing index was calculated as shown in Fig. 2A. Colored dots represent significantly differentially edited dsRNAs. Orange dots, 3’ UTR dsRNAs. Green dots, intergenic dsRNAs. **b**, Fold-enrichment of edited dsRNAs in TRIM65-pulldown. The same edited dsRNAs shown in (**a**) were plotted. **c**, **d**, Enrichment analysis of TRIM65-pulldown signals in differentially edited (**c**) versus non-differentially edited dsRNAs (**d**). We divided each edited dsRNA sequence ±50nt into 60 equal bins, and within each bin, we calculated the fold-enrichment of TRIM65-pulldown over input for MDA5 WT and MDA5 KO cells. 5’, 5’ end of the dsRNA sequence; 3’, 3’ end of the dsRNA sequence. **e**, Venn diagram showing the overlap among differentially edited dsRNAs (DEDs), non-differentially edited dsRNAs (non-DEDs), TRIM65 pull-down RNAs in MDA5 WT cells (Pulldown-MDA5), and TRIM65 pull-down RNAs in MDA5 KO cells (Pulldown-bkg). **f**, Comparison of DED vs non-DED enrichment in TRIM65 pull-down experiment. DEDs show strong enrichment, whereas non-DEDs exhibit minimal enrichment, indicating that differentially edited dsRNAs preferentially act as MDA5-associated ligands in the TRIM65 pull-down.

Consistent with this specificity, among the 1,460 non-DED dsRNAs present in both control and DDX3X-depleted cells (Fig.4a, gray dots), only 13 were detected specifically in TRIM65-purified MDA5 pulldowns, representing <0.9% of the total (Fig.4a,b,e,f). This is in line with prior estimates that only a small minority of edited dsRNAs are competent to activate MDA5^13,24^. In addition, 212 dsRNAs were detected in TRIM65 pulldowns but not identified by RNA editing analysis and were absent from TRIM65 pulldowns in MDA5 knockout cells (Fig.4e). These dsRNAs included Alu elements with relatively low editing frequencies as well as other repetitive elements, such as LINEs, simple repeats, DNA transposons, LTRs, and centromeric repeats (Fig.3e), suggesting the existence of additional classes of MDA5-associated dsRNAs that are not captured by differential editing analysis alone.

### DDX3X and ADAR1 cooperatively suppress inverted-repeat dsRNA formation in 3′UTRs

To examine whether RNA helicases that suppress ISG responses and ADAR1 act on overlapping RNA substrates, we analyzed publicly available sequencing datasets from prior studies that profiled DDX3X, ADAR1, and DHX9 binding interactions with RNA^20,50,51^. Metagene-analysis of these datasets revealed distinct but partially overlapping transcript positional preferences. DDX3X showed predominant enrichment in 5′UTRs as previously described^50,52^ with additional, albeit weaker, enrichment in 3′UTRs. Both ADAR1 and DHX9 were primarily enriched in 3′UTRs, with detectable but reduced association in 5′UTRs (Fig.5a). Consistent with these positional biases, correlation analysis of RNA-seq, RIP-seq, and CLIP-seq datasets revealed positive correlations among DDX3X-, ADAR1-, and dsRNA-associated signals within 3′UTRs, whereas correlations were markedly weaker within 5′UTRs (Fig.5b). This positional link between ADAR1 and DDX3X within 3’ UTRs was also stronger than that of ADAR1 and DHX9 (Fig.5b; Pearson correlation coefficient 0.55 versus 0.38, respectively). These findings indicate that, despite differing global binding preferences, DDX3X and ADAR1 frequently converge on shared 3′UTR regions across different cell types, consistent with overlapping substrate pools.

**Fig. 5.**
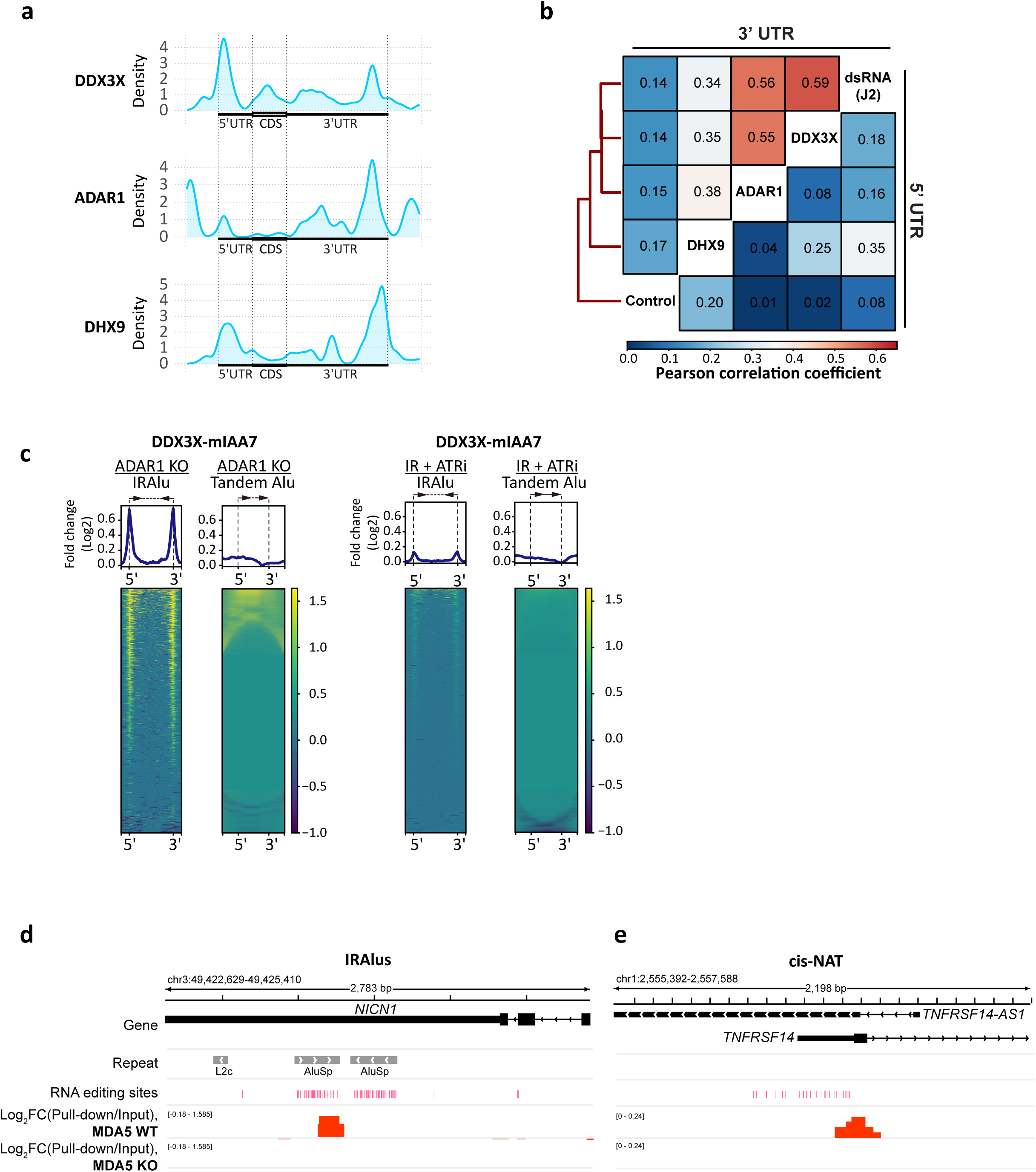
Analysis of shared DDX3X and ADAR1 substrate dsRNAs present in MDA5 filaments. **a**, Metagene analysis of publicly available data showing common binding enrichment of DDX3X, ADAR1, and DHX9 for 3′UTRs. **b**, Correlation analysis of control RNA-seq (Control), dsRNA (J2) RIP-seq (dsRNA (J2)), DDX3X eCLIP-seq (DDX3X), ADAR1 CLIP-seq (ADAR1), and DHX9 RIP-seq (DHX9) signals within 3′UTRs and 5’UTRs. **c**, Heatmaps showing enrichment of inverted-repeat Alu (IRAlu) dsRNAs following co-depletion of DDX3X and ADAR1(DDX3X-mIAA7 + ADAR1 KO) or loss of DDX3X combined with IR + ATRi (DDX3X-mIAA7 + IR +ATRi). **d**, Genome browser view of the *NICN1* locus illustrating IRAlu elements within the 3′UTR, RNA editing sites, and enrichment of dsRNAs recovered by TRIM65 pull-down in MDA5 wild-type but not MDA5 knockout cells. **e**, Genome browser view of the *TNFRSF14:TNFRSF14-AS1* locus showing sense-antisense *cis*-natural antisense transcript (*cis*-NAT)-derived dsRNAs, RNA editing sites, and MDA5-dependent enrichment in TRIM65 pull-downs.

We next asked whether simultaneous loss of DDX3X and ADAR1 function alters the structural organization of dsRNAs arising from these shared substrates. Notably, inverted repeat Alu elements (IRAlus) that form dsRNAs but not tandem Alus were highly enriched after ADAR1 and DDX3X co-depletion (Fig.5c). In comparison, there was a reduced level of overlapping IRAlus enriched in the DDX3X-mIAA7 cells that were treated with IR + ATRi (Fig.5c). This is consistent with a model that ADAR1 limits the pool of dsRNAs that accumulate in DDX3X deficient cells from entering active MDA5 filaments. Analysis of the DEDs and non-DEDs revealed that more than 90% of dsRNA in both groups were comprised of IRAlus (Extended Data Fig.3b,c). However, the marked differences in their enrichment within active MDA5 filaments prompted a closer examination of DED dsRNA structure and editing patterns. We observed canonical MDA5 substrates derived from IRAlus within genes such as *NICN1*, *DES1*, *DHFR*, *GNINS1*, and *MAVS*, which are known to activate wild-type MDA5 *in vitro*^28^ (Fig.5d). In addition, we identified several examples of sense-antisense dsRNAs arising from *cis*-NATs dsRNAs produced from *TNFRSF14* and *DDTL* loci (Fig.5e), revealing endogenous dsRNA sources that require dual regulation by DDX3X and ADAR1 to suppress aberrant MDA5 activation.

We next asked whether the immunogenic dsRNAs identified in DDX3X depleted RPE-1 cells would be observed across diverse cell and tissue types that harbor *DDX3X* loss of function mutations. Somatic *DDX3X* mutations are common in several cancer types. To assess whether similar alterations occur in human cancers, we analyzed tumors harboring *DDX3X* unwinding-deficient mutations. We observed widespread tumor-specific shifts in RNA editing in both lung adenocarcinoma (LUAD) and uterine and endometrial cancer (UCEC) harboring loss of function RNA helicase *DDX3X* missense alleles (T275M) and (R534H), respectively (Fig.6a,b)^40,53^. The most significantly altered sites showed higher RNA editing level in the tumor relative to matched normal tissue^36,54^. Importantly, 73% of the dsRNAs more highly edited in tumors were present in the DED group defined in RPE-1 cells. In contrast, only 3 dsRNAs more highly edited in matched normal tissues were present in the DEDs. Together, these findings demonstrate that *DDX3X* loss or mutation promotes increased RNA editing of a conserved set of potential MDA5 ligands across cellular, *in vivo*, and disease contexts.

**Fig. 6.**
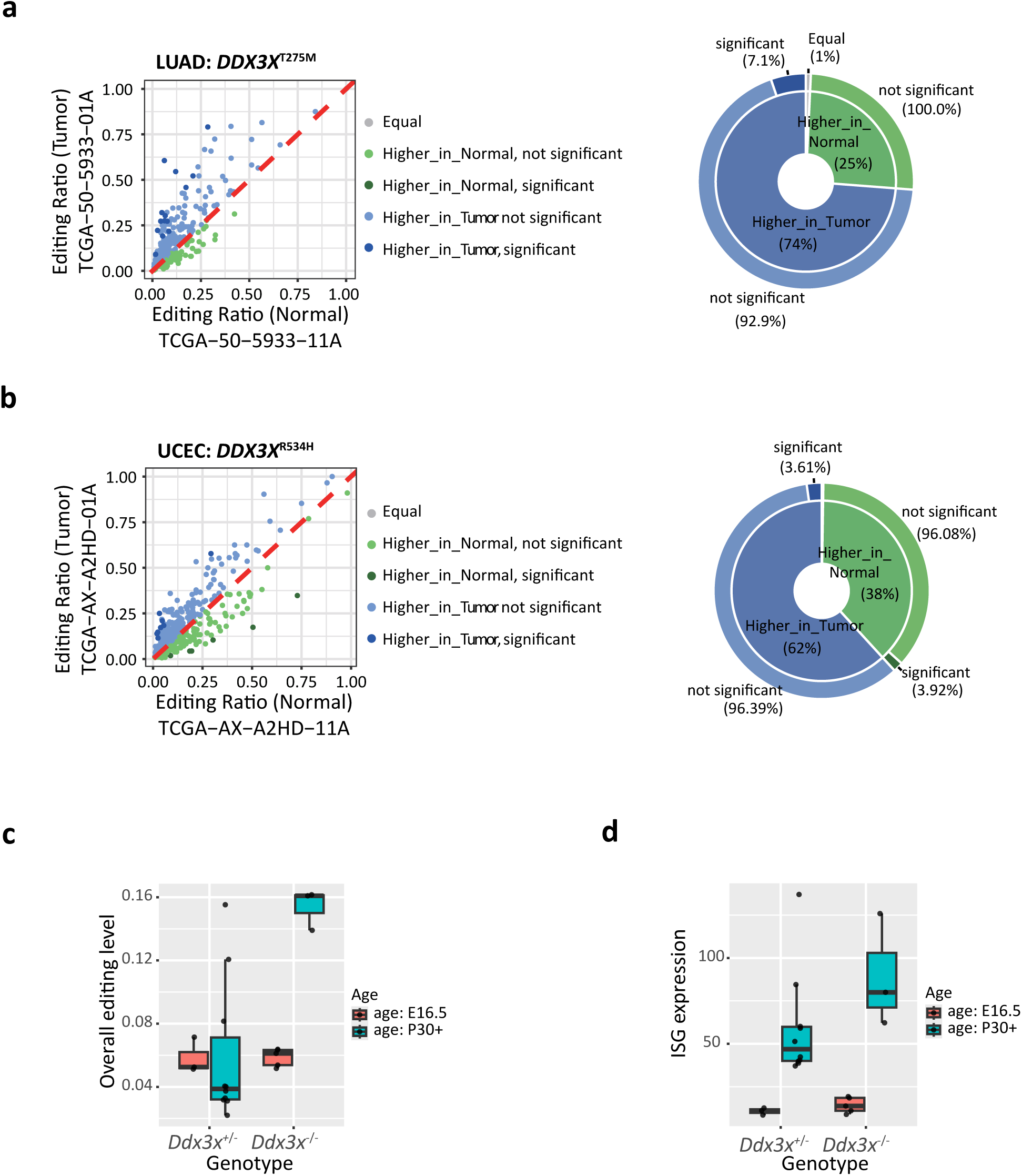
DDX3X loss-of-function mutations lead to elevated editing on the identified MDA5 ligand dsRNAs. **a**, Left: RNA editing differences between tumor and matched solid normal samples in a lung adenocarcinoma (LUAD) patient harboring the *DDX3X*^T275M^ somatic mutation. Right: Pie chart depicting the proportion of editing sites exhibiting significant differential editing. **b**, Left: RNA editing differences between tumor and matched solid normal samples in a uterine corpus endometrial carcinoma (UCEC) patient harboring the *DDX3X*^R534H^ somatic mutation. Right: Pie chart depicting the proportion of editing sites exhibiting significant differential editing. **c**, Overall RNA editing levels in *Ddx3x*^+/-^ and *Ddx3x*^-/-^ mice at embryonic day 16.5 (E16.5) and postnatal day 30 (P30). **d**, Interferon-stimulated gene (ISG) expression levels were calculated using a standard ISG score gene list (*Isg15, Mx1, Ifit3, Ifit1, Oas2, Usp18, Rsad2, Ifi44*) in *Ddx3x*^+/-^and *Ddx3x*^-/-^ mice at E16.5 and P30.

*DDX3X* mutations are common in human medulloblastoma and targeted deletion *Ddx3x* in the brain promotes medulloblastoma in mice^55^. We analyzed RNA seq data from *Ddx3x*^+/-^ and *Ddx3x*^-/-^ mice at embryonic day 16.5 (E16.5) and at 30 days post-development^55^. In further agreement with Ddx3x being responsible for suppressing immunogenic dsRNAs, *Ddx3x*^-/-^ mice exhibited markedly increased global RNA editing and moderately elevated ISG expression at P30 (Fig.6c,d). Interestingly, Ddx3x-dependent editing and ISG elevation was not observed at E16.5, indicating a developmental stage-dependent requirement for Ddx3x in constraining immunogenic dsRNA formation and suppressing aberrant immune activation^55^.

### Dual dsRNA substrates exhibit higher repeat complementarity and lower free energy

DEDs and non-DEDs were predominantly derived from Alu elements. We stratified the TRIM65 pull-down peaks into four groups: DEDs, non-DEDs, TRIM65 pull-down excluding DEDs and non-DEDs, and MDA5 KO background (Fig.4e). DEDs and non-DEDs exhibited higher overall editing levels than dsRNA detected exclusively in the TRIM65 pull-down or MDA5 KO background (Fig.7a). Moreover, RNA editing of those shared 246 dsRNAs was approximately 2-fold higher in DDX3X-depleted cells than in control cells (Fig.4e and Fig.7a). By contrast, the 46 dsRNAs absent from the TRIM65 pull-down displayed markedly lower RNA editing levels (Fig.4e and Fig.7a). Analysis of RNA editing patterns further revealed that distances between inverted repeat sequences were significantly increased upon loss of DDX3X (Fig.7b). In DDX3X-depleted cells, the average distances between editing clusters in DDX3X depleted cells were larger, and 11.3% of the IRAlus showed intervening distances greater than 1 kb. The non-DEDs showed less than 1.9% with intervening distances of more than 1 kb. These data indicate that DEDs preferentially arise from inverted repeat elements separated by longer genomic intervals.

**Fig. 7:**
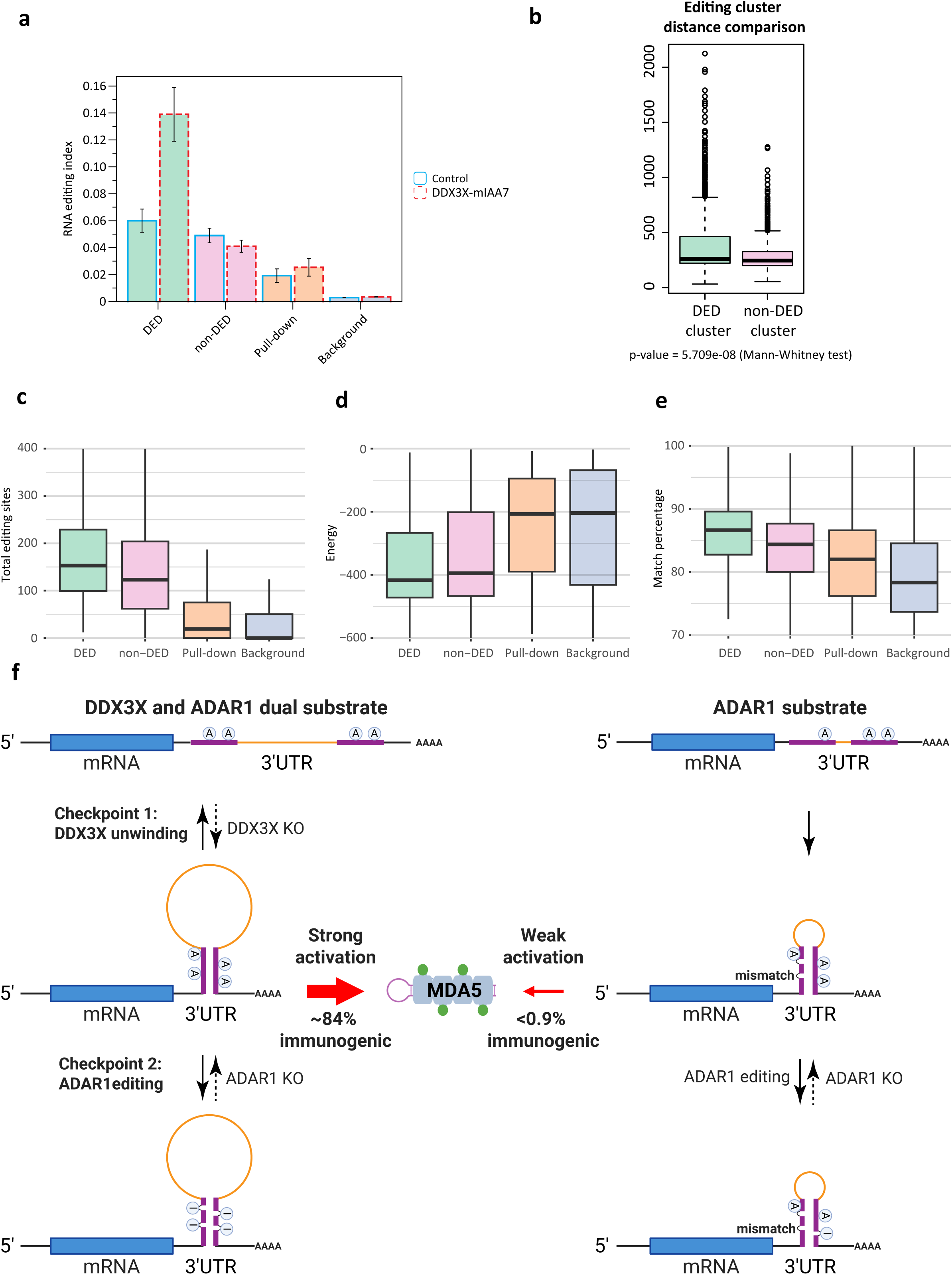
Characteristics of highly immunogenic DDX3X-ADAR1 dual substrate dsRNAs. **a**, Comparison of RNA-editing levels across differentially edited dsRNAs (DEDs), non-differentially edited dsRNAs (non-DEDs), TRIM65 pull-down RNAs (Pull-down), and background RNAs (Background) in control and DDX3X-depleted (DDX3X-mIAA7) cells. **b**, Comparison of RNA editing cluster distances for dual DDX3X-ADAR1 substrates (DEDs) and non-DEDs; p value determined by Mann–Whitney test. **c-e**, Structural and sequences features of dsRNA classes, including RNA editing levels (**c**), predicted minimum free energy (**d**), and sequence match percentage between paired repeats (**e**) for DEDs, non-DEDs, pull-down RNAs, and background RNAs. **f**, Model illustrating dual checkpoint regulation of 3′UTR dsRNAs by DDX3X-mediated unwinding and ADAR1-mediated editing. Left: Dual DDX3X–ADAR1 substrates consist of inverted Alu repeats with high sequence complementarity and longer inter-repeat distances, forming stable dsRNAs that elicit strong MDA5 activation when both checkpoints are lost. Right: ADAR1-only substrates contain inverted Alu repeats with greater sequence mismatches and shorter inter-repeat distances, producing dsRNAs that have limited potential for MDA5 activation.

Consistent with these structural features, DEDs harbored a greater number of editing sites, exhibited lower predicted minimum free energy, and displayed fewer sequence mismatches between paired repeats compared with non-DEDs, pull-down RNAs, and background RNAs (Fig.7c-e). Collectively, these findings support a model in which highly stable dsRNAs with long inter-repeat distances and minimal mismatches represent a class of potent innate immune ligands that require coordinated suppression by both DDX3X-mediated unwinding and ADAR1-mediated editing (Fig.7f). While such dsRNAs are efficiently destabilized by DDX3X in wild-type cells (Checkpoint 1), their increased stability and complementarity render them particularly effective ADAR1 substrates upon DDX3X depletion (Checkpoint 2), thereby necessitating dual regulation to prevent excessive innate immune activation.

## Discussion

Hyperactive MDA5 signaling produces severe inflammation and a lethal form of autoimmunity. These genetic observations provide compelling evidence that endogenous immunogenic dsRNAs are produced as a byproduct of normal transcriptional processes in unperturbed cells. The nature of the putative endogenous RNA immunogens has remained enigmatic as have the quality control mechanisms beyond A-to-I editing that suppress their formation. Several barriers exist that hinder the identification of sources of endogenous immunogenic dsRNAs. Although MDA5 is exclusively activated by dsRNA ligands, it binds with high-affinity to dsRNA, dsDNA, and ssRNA^1^. Consequently, direct purification of MDA5 monomers leads to an excessive background of ssRNA, making it difficult to identify the activating immunogenic dsRNAs. An *in vitro* RNase protection assay suggested hyperactive Aicardi-Goutières Syndrome associated MDA5 mutants binding to IRAlu sequences to promote autoimmune disease^28^. Additional evidence supports IRAlu activation of dsRNA pattern recognition receptors in cancer^56,57^. Compared to the hyperactive AGS-associated MDA5 mutant, wild type MDA5 is more sensitive to A-to-I edits present in double-stranded regions of inverted repeat RNAs, which limits inappropriate inflammatory signals^27,28^. In accordance, less than 2% of ADAR1 edited IRAlus are predicted to activate MDA5^58^, indicating that most IRAlus are insufficient to activate wildtype MDA5 and additional determinants for dsRNA immunogenicity are necessary.

In this study, we leverage the specificity of the TRIM65 E3 ubiquitin ligase for oligomeric MDA5 to purify the active MDA5 filament and sequence its dsRNA cargo. Coupled with CRISPR screening approaches and analyses of A-to-I editing profiles, we determine that suppression of a class of highly active MDA5 ligands requires cooperative DDX3X RNA helicase and ADAR1 base editing activities. More than 80% of the differentially edited RNAs/DEDs were present in purified MDA5 filaments from cells that lack DDX3X and ADAR1. Greater than 90% of these dual substrate dsRNAs were IRAlu elements embedded within RNA Polymerase II transcripts that contain high repeat complementarity, longer distances between inverted repeat elements, and lower predicted free energy. These immunogenic 3’-UTR associated IRAlu elements demonstrate shared association with DDX3X and ADAR1 across cell types (Fig.5a,b), consistent with their designation as dual substrates (Fig.7f).

While the structural basis for shared substrates between DDX3X and ADAR1 is yet to be defined, several features of the DED RNAs may offer insights. DDX3X unwinding occurs preferentially on substrates with 3’-single stranded RNA overhangs^59–61^. IRAlus embedded within DED RNAs were present close to the terminus of the 3’-UTR, which predicts a 3’-ssRNA overhang that is ideally suited to DDX3X recognition and unwinding of proximal dsRNA in paired IRAlus. The presence of *cis*-NAT RNAs as dual substrates present in MDA5 filaments, suggests that additional classes of MDA5 ligands also require suppression by combined RNA helicase and base editing activities. It is important to note that while ADAR1 edited dsRNAs in DDX3X-expressing cells were also predominantly IRAlus, less than 1% of them are associated with MDA5. This demonstrates that DDX3X specifically suppresses the formation of a class of highly active MDA5 ligands, whereas most IRAlus are not immunogenic and are acted on solely by ADAR1.

The canonical function of DDX3X has largely been ascribed to its role in translation initiation, where it resolves highly structured 5’ untranslated regions (5’UTRs) to facilitate ribosome scanning.

Our findings unveil a critical, spatially distinct function for DDX3X at the 3’ ends of transcripts. We suggest that DDX3X is essential for unwinding highly complementary inverted repeat Alu (IRAlu) elements embedded predominantly within 3’UTRs (Fig.7f). This shifts the paradigm of DDX3X from primarily a translational regulator to a vital guardian of innate immune homeostasis. Because DDX3X preferentially unwinds substrates with 3’-single-stranded RNA overhangs, the architecture of 3’UTRs terminating in poly(A) tails likely provides an optimal structural context for its helicase activity. Consequently, the cooperative action of DDX3X and ADAR1 on these 3’UTR-localized IRAlus highlights a specialized, translation-independent quality control mechanism that prevents endogenous transcripts from adopting immunogenic, viral-like dsRNA conformations (Fig.7f).

Hypomorphic mutations in ADAR1 are causative for Aicardi-Goutières, a pediatric lupus like syndrome characterized by autoimmunity and brain calcifications^62,63^. DDX3X is an X-linked RNA helicase that escapes X-inactivation in females^64^. DDX3Y is the Y chromosome encoded homolog of DDX3X but harbors several features that make it less active^65,66^. Germline loss of function mutations in DDX3X are lethal in males and *de novo* heterozygous DDX3X mutations in females produce a neurodevelopmental syndrome that entails intellectual disability autism, epilepsy, and gross motor dysfunction^53,67^. Elevated inflammation and autoimmunity have not been reported in DDX3X mutant patients^53,67^. There are several possible explanations for phenotypic differences between ADAR1 and DDX3X mutation carriers. Our findings suggest that ADAR1 editing of a specific class of IRAlu RNAs in DDX3X deficient cells would limit immunogenic dsRNA formation. Additionally, ADAR1 associated syndromes arise from biallelic hypomorphic mutations, whereas DDX3X mutations typically manifest in the heterozygous state and do not recapitulate the degree of DDX3X deficiency present in our experiments. DDX3X RNA helicase has also been reported to promote translation of specific RNAs by unwinding secondary structures in 5’-UTR regions and thus has additional roles beyond suppressing MDA5 activation that may contribute to developmental syndromes in germline heterozygous carriers.

Somatic inactivating DDX3X missense mutations are frequent in several malignancies, including medulloblastoma and lymphomas^68,69^. DDX3X mutations are also common in virus associated malignancies^70^. While it is unclear how this provides a selective advantage in specific cancers, emerging evidence indicates that chronic inflammatory signaling from hyperactive pattern recognition receptor signaling can promote tumor formation and metastasis^71,72^. In agreement, diverse human tumors that harbor missense mutations in the DDX3X helicase domain also displayed higher editing, with the majority of hyper-edited RNAs overlapping with the DDX3X substrates defined in our study (Fig.6a,b). Analysis of RNA-seq data in DDX3X mutant mouse medulloblastomas also demonstrated both higher editing indices and ISG levels (Fig.6c,d). This provides evidence that DDX3X mutant cancers display increased ISG levels and that differentially edited immunogenic dsRNAs defined in our study are dually regulated by DDX3X and ADAR1 across diverse cell and tissue types. These findings also suggest therapeutic possibilities and predictive biomarkers derived from DDX3X mutation. Complete ADAR1 deficiency is rapidly lethal in adult mice, whereas inhibition of its base editing enzymatic activity is better tolerated. Indeed, ADAR1 inhibitors are entering clinical trials for acute myeloid leukemia (NCT07250646). ADAR1 loss of function overcame immune checkpoint inhibition resistance in syngeneic mouse models^73^ and ADAR1 was required for viability in a subset of cancer cells with increased ISG expression^74^. Our findings raise the possibility that ADAR1 inhibition would produce lethal levels of inflammation in DDX3X mutated cancers that would hinder tumor growth and perhaps stimulate productive responses to immune checkpoint inhibition immunotherapies. This justifies a rationale to investigate whether elevated ADAR1 editing of the identified highly immunogenic dsRNAs defined in this study could represent a specific biomarker for the productive use of ADAR1 inhibitors.

## Methods

### Cell lines and culture conditions

The hTERT RPE-1 cell line was obtained from ATCC. The HEK293T cell line and hTERT RPE-1-derived cell lines, including hTERT RPE-1 p53 KO and hTERT RPE-1 p53 KO DDX3X-mIAA7, were cultured in DMEM with 10% bovine calf serum (Hyclone). The hTERT RPE-1 p53 KO DDX3X-mIAA7 cell lines with inducible wild type DDX3X or mutated DDX3X expression were maintained in DMEM supplemented with 10% Tet-free fetal bovine serum (Omega Scientific). All cell lines were maintained with penicillin and streptomycin (Thermo Fisher Scientific) in a humidified incubator with 5% CO_2_ at 37°C.

### Plasmid construction

LentiGuide-puro vectors targeting different genes were generated by inserting targeting duplex oligos into LentiGuide-puro vector (gift from Dr Feng Zhang, Addgene #52963) at the BsmBI site. pSpCas9(BB)-2A-GFP(PX458) vectors targeting different genes were generated by inserting targeting duplex oligos into pSpCas9(BB)-2A-GFP(PX458) vector (gift from Dr Feng Zhang, Addgene #48138) at BbsI site. Oligos for gRNAs are listed in Supplementary Table 1. Donor plasmids of pGL3-DDX3X-mIAA7-3XFLAG-HA_AtAFB2-weak NLS-Blast/Neo were generated by HIFI assembly using the pGL3 backbone, AtAFB2-weak NLS (both amplified from pSH-EFIRES-P-AtAFB2-mCherry-weak NLS, gift from Dr Elina Ikonen, Addgene #129717), DDX3X homologous arms (amplified from genomic DNA of hTERT RPE-1 p53 KO cells), and synthesized gene fragments encoding mIAA7-3XFLAG-HA and BSD/Neo (IDT DNA). pCW57-GFP-P2A-DDX3X vector was generated by inserting GFP (using pmGFP-ADAR1-p150 as template, gift from Dr Kumiko Ui-Tei, Addgene #117927) and DDX3X (using pHAGE-DDX3X as template, gift from Dr Keneth Scott, Addgene #116730) into the same reading frame within all-in-one Doxycycline-inducible pCW57-MSC1-P2A-MCS2 (neo) vector (gift from Dr Adam Karpf, Addgene #89180). DDX3X AAA, G302V, and G325E mutations were generated using pDONR221-DDX3X as previously described^75^. DDX3X R534H and R475G mutations were generated as previously described^40^. All pCW57-GFP-P2A-DDX3X mutants were created by replacing wild-type DDX3X in pCW57-GFP-P2A-DDX3X with PCR amplified DDX3X mutant fragments using HIFI assembly (NEB). pcDNA3-V5-SBP-HA-TRIM65 (CC-PSpry) was generated by inserting a synthesized oligo encoding V5-SBP-(GGGGS)3 linker into the pcDNA3-HA-TRIM65 (CC-PSpry) at N-terminal of HA-TRIM65.

### mIAA7 degron cells

2.25 million hTERT RPE-1 p53 KO cells were seeded in a 15 cm dish one day before transfection. 1 hour before transfection, cells were replaced with 25 ml of fresh medium. 3.5 μg CRISPR-Cas9 all-in-one plasmid pSpCas9(BB)-2A-GFP(PX458) plasmid, targeting the vicinity of the stop codon, along with 2 μg each of two donor plasmids (neo/blast) providing templates for mIAA7 tag knock-in and AtAFB2 expression for auxin-induced degradation of mIAA7-tagged proteins, were transfected into cells using 10.5 ul Avalanche-Omni transfection reagent. 4 hours post transfection, cells were replaced with 25 ml of fresh medium. 2 days post-transfection, GFP-positive cells were sorted and cultured for 10-14 days followed by selection with 400 μg/ml G418 (Corning) and 10 μg/ml blasticidin (Invivogen) for at least 14 days. After withdrawing G418 and blasticidin, cells might require an additional 7-14 days to recover normal protein expression prior to western blot analysis of the knock-in.

The target protein was tagged with a (GGGGS)_3_ linker, an mIAA7 degron tag, a 3xFLAG tag, and an HA tag, adding approximately 13 kDa to its molecular weight. This allowed differentiation between knock-in and non-knock-in proteins by western blot. Knock-in efficiency was estimated by comparing parental cells with knock-in cells, including a 10 % loading for both, via western blot. The positive knock-in population can be determined by probing the 3xFLAG tag using anti-FLAG M2 antibody (Millipore Sigma, F1804-200UG, which is more sensitive than the HA antibody) or the HA tag using anti-HA antibody (Genscript, A01244-100, if a FLAG-tagged protein is already present in cells). This can be assessed by FACS for high-expression proteins, or by immunofluorescence staining for proteins with distinct staining patterns. Depletion efficiency was validated by comparing parental cells with knock-in cells upon treatment with 1 mM auxin for different durations. In most cases, a 4-hour treatment achieved over 90% depletion.

Due to the high efficiency of homozygous knock-in (>95%) observed across the tested cell lines and genes, single-clone screening was not performed. Instead, bulk polyclonal cells were used to minimize clonal variation.

### Plasmid-based CRISPR–Cas9 knockout

hTERT RPE-1 p53 KO cell line was generated by transfecting pSpCas9(BB)-2A-GFP(PX458) vector targeting p53 into hTERT RPE-1 cell line (ATCC). 2 days post transfection, GFP positive cells were sorted and seeded at one cell per well in 96-well plate, and single clones were identified and marked under microscope 4 days later. Single clones were expanded, passaged, and validated by western blot.

### Lentiviral CRISPR–Cas9 knockout

Lentiviruses of LentiCas9-mCherry (gift from Dr Junwei Shi, University of Pennsylvania), LentiCas9-blast and LentiGuide-Puro (gifts from Dr Feng Zhang, Addgene #52962, #52963) were prepared and concentrated as previously described^76^. Cells were infected with lentiviruses overnight in the presence of 8 μg/ml polybrene. Cells infected with LentiCas9-Blast were subjected to 10 μg/ml blasticidin (Invivogen) until control cells without infection were all dead. Cells infected with LentiCas9-mCherry were sorted using an FACSAria sorter (BD). Knockouts were created by infecting Cas9-expressing cells with lentivirus produced using lentiGuide-Puro. To screen single colonies, cells were seeded one cell per well in 96-well plate and single clones were identified and marked under microscope 4 days later. Single clones were expanded, passaged, and validated by western blot. Otherwise, cells were cultured for 4 days before experimental treatment. Knockouts were validated by western blot using respective antibodies.

### Irradiation and drug treatments

Cells were seeded to allow sufficient space for proliferation, achieving a density of 40%-50% at the time of collection. Irradiation was performed using a Cs-137 Gammacell irradiator (Nordion) with a dose rate of ∼0.667 Gy/min. ATR inhibitor (ATRi) was administrated 1 h prior to irradiation to inhibit ATR activity. Doxycline was applied 1 day before treatment to induce wild type or mutated DDX3X expression. Unless otherwise indicated, auxin was applied 3 hours prior to treatment to induce depletion of DDX3X-mIAA7. Drugs were maintained in medium throughout the experiment, and medium with or without drugs was refreshed every 2 days unless noted otherwise. The concentrations of drugs were used as follows: ATR inhibitor (ATRi; VE-821, 2.5 μM; Selleck Chemical), Doxycycline (DOX; 2 μg/ml; Sigma), and Auxin (3-Indoleacetic acid, 1 mM, Sigma).

### Array-based lentiviral CRISPR–Cas9 screening

Three individual sgRNAs targeting different functional domains for each gene were designed, cloned into lentiGuide-Puro vector, and packaged into lentiviral sgRNAs. Three individual lentiviral sgRNAs for each gene were pooled in equal volumes, aliquoted to 0.5 ml per well in a 1 ml deep 96-well PP plate, sterile (USA Scientific, 1896-1110), sealed with autoclaved AlumaSeal 96 Sealing Foil (Excel Scientific, F-96-100), and stored in a −80°C deep freezer. In a pilot experiment, control lentiviral sgRNAs and positive lentiviral sgRNAs (using RIG-I sgRNAs. Lentiviral sgRNAs of one gene in each well) were used at different dilutions to infect hTERT RPE-1 p53 KO cells with Cas9-blast expression in 24-well plates in the presence of 8 μg/ml polybrene. The next day, cells were passaged to 6-well plates and cultured for another two days. 3 days post-infection, cells were seeded into new 6-well plates. 4 days post-infection, the medium was replaced with fresh medium containing 2.5 μM ATR inhibitor (VE-821) for 1 hour, followed by 10 Gy irradiation. Cells were cultured for 3 days with medium replacement containing ATR inhibitor every 2 days. Cells were collected for western blot analysis to determine the minimum viral dilution required for efficient knockout. Once the optimal lentiviral dilution was determined, a new plate of lentiviral sgRNAs was thawed for screening using method as described above, except cells were collected for RT-qPCR analysis.

### RT-qPCR

Total RNA was extracted using the Direct-zol RNA Miniprep kit (Zymo, R2050) with on-column DNase I digestion (Qiagen, 79254). 1 μg RNA was used for reverse transcription with High Capacity cDNA Reverse Transcription Kit (Thermo Scientific, 4368814). Synthesized cDNAs were diluted 20-fold for qPCR analysis using Applied Biosystems Power SYBR Green PCR Master Mix (Thermo Scientific, 4367659) on an ABI 7900HT system (Thermo Scientific). Oligos for RT-qPCR are listed in Supplementary Table 1.

### Western blot

Western blot was performed as previously described^29^. In brief, cells were collected by trypsinization, washed in PBS and lysed in NETN buffer (150 mM NaCl, 1% NP-40 alternative, 50 mM Tris pH 7.4) with turbo nuclease (Accelagen) in the presence of 1 mM MgCl_2_, protease inhibitor cocktail (Roche) and Halt phosphatase inhibitor cocktail (Thermo Scientific) for 30 min on ice. Protein was vortex omitting centrifuge prior to quantification using Pierce BCA protein assay (Thermo Scientific). Equal amount of protein was boiled with 4x NuPAGE LDS Sample Buffer (Thermo Scientific, NP0007) containing 10% 2-mercaptoethanol at 94°C for 5 minutes, then separated on Bolt 4-12 % Bis-Tris Plus gels (Thermo Scientific) using MOPS/MES buffer and transferred to 0.2 µM Amersham Protran Premium nitrocellulose western blot membrane at 350 mA for 1 hour and 20 minutes in ice cold transfer buffer (20 % methanol, 191 mM glycine, 25 mM tris-base, 0.1% SDS, pH 8.3). Membrane was blocked in 3% non-fat milk in TBST (0.1% Tween 20) at room temperature for 10 minutes and incubated in primary antibody diluted in 1% BSA in PBS overnight at 4°C. Membrane were washed in TBST for 15 min and incubated in secondary antibody (Amersham ECL HRP, GE) diluted in 3% non-fat milk in TBST at room temperature for 2 h. Blots were developed using Immobilon Forte Western HRP substrate (Millipore Sigma).

### Silver staining

Silver staining was performed using the SilverQuest Silver Staining Kit (Thermo Scientific, LC6070) according to the manufacturer’s instructions. Briefly, purified TRIM65 was boiled with 4x NuPAGE LDS Sample Buffer (Thermo Scientific, NP0007) containing 10% 2-mercaptoethanol at 94°C for 5 minutes and then separated on Bolt 4-12 % Bis-Tris Plus gels (Thermo Scientific) using MOPS buffer. After electrophoresis, the gel was briefly rinsed with ultrapure water and fixed in a solution containing 40% ethanol and 10% acetic acid (prepared with ultra-pure water) for 20 minutes. The gel was then washed in 30% ethanol for 10 minutes, followed by incubation with sensitizing solution for 10 minutes. Next, the gel was washed with 30% ethanol for 10 minutes, followed by ultrapure water for 10 minutes, before incubation in staining solution for 15 minutes. The stained gel was briefly rinsed with ultrapure water for 20-60 seconds, then developed in developing solution until the desired band intensity was reached. Finally, stopper solution was added, and the gel was incubated for 10 minutes to complete the staining process.

### Immunofluorescence

Cells were seeded onto coverslips (Electron Microscopy Sciences, 72230-01) in 24-well plates one day before treatment and fixed with 3% paraformaldehyde (PFA) at room temperature for 10 minutes. For cytoplasmic dsRNA or ISG56 staining, all the staining and washing procedures were performed on ice or at 4°C. Cells were washed with PBS, permeabilized with 0.1% Saponin in PBS for 1 min, and processed for immunostaining using the indicated antibodies with 0.1% Saponin in PBS. Stained cells were finally washed with PBS and fixed in 1x Stabilizing Fixative (BD) at room temperature for 10 minutes. Coverslips were mounted in VECTASHIELD PLUS Antifade Mounting Medium with DAPI (Vector Laboratories). Images were taken using a Nikon Eclips 80i microscope with a Coolsnap Myo camera (Photometrics) and Nikon NIS-Elements software. Images and fluorescence intensity measurement were prepared using FIJI (NIH). Statistical graphs were prepared using Prism (Graphpad).

### Flow cytometry

Cells were dissociated into single cells with trypsin from one six-well, washed once in PBS and fixed with 3% paraformaldehyde (PFA) at room temperature for 10 minutes. For cytoplasmic dsRNA or ISG56 staining, all the staining and washing procedures were performed on ice or at 4°C. Cells were washed with PBS, permeabilized with 0.1% Saponin in PBS for 1 min, and processed for immunostaining using the indicated antibodies with 0.1% Saponin in PBS with occasional resuspension using pipetting. Stained cells were finally washed and resuspended with 1% BSA in PBS. Flow cytometry was performed on a MACSQuant Analyzer with a precooled Chill 96 Rack (Miltenyi Biotec) and analyzed with FlowJo software.

### Cytosolic RNA extraction for RNA editing analysis

All procedures were performed in RNase-free environment at 4°C or on ice. Cytosolic extracts were prepared using digitonin extraction as previously described^44^. Briefly, cells were resuspended in a 1.7 ml EP tube with buffer containing 150 mM NaCl, 50 mM HEPES pH 7.4, and 25 μg/ml digitonin (Sigma, D141). The resuspended cells were incubated with end-over-end rotation for 10 minutes to allow selective plasma membrane permeabilization, followed by centrifuge at 980g for 3 minutes three times to pellet intact cells. The cytosolic supernatants were transferred to new tubes and centrifuged at 17,000g for 10 minutes to pellet any remaining cellular debris, yielding cytosolic preparations free of nuclear, mitochondrial, and endoplasmic reticulum contamination. The cytosolic RNA extract was then processed using Direct-zol RNA Miniprep kit (Zymo, R2050) with on-column DNase I digestion (Qiagen, 79254) for RNA extraction.

### TRIM65 purification

Please also see (Chen, Kim, et al. Biorxiv 2026) for additional description. HEK 293T cells were seeded in six 15 cm dishes at 12 million cells per dish one day before transfection. One hour before transfection, cells were replaced with 25 ml of fresh medium. For each dish, 30 μg of pcDNA3-V5-SBP-HA-TRIM65 was transfected into cells using 42 μl Avalanche-Omni transfection reagent. Cells were cultured for three days with daily medium replacement.

All subsequent procedures were performed in RNase-free environment at 4°C or on ice unless otherwise indicated. Cells from each dish were washed twice with 7 ml PBS, scraped into 4 ml PBS, pooled, and centrifuge to pellet. The cell pellet (∼2 ml) was resuspended in 18 ml modified hypotonic buffer (10 mM Tris pH 7.5, 10 mM KCl, 1.5 mM MgCl_2_, 5% glycerol, fresh 1 mM TCEP and protease inhibitor cocktail), then aliquoted into two 50 ml LoBind tubes (Eppendorf, 0030122240) with 9 ml each and incubated for 15 minutes. Cells were lysed by gently passing the resuspension through a 25G needle fitted to a 10 ml syringe 7 times, avoiding bubbles. The cell lysates were aliquoted into 1.7 ml EP tubes and sequentially centrifuged at 1,000 g, 5,000 g and 18,000 g for 5 minutes each. The resulting cytoplasmic supernatant (∼10 ml) was carefully pooled into a 50 ml LoBind Eppendorf tube, mixed with an equal volume (10 ml) of modified RBB buffer (20 mM HEPES pH 7.5, 1. 5mM MgCl_2_, 1 M NaCl, 5% glycerol, 0.02% NP-40 and fresh 1 mM TCEP, and protease inhibitor cocktail), and precleared with 120 ul of TrueBlot Protein G magnetic beads (Rockland, PG00-18-2) by gentle rotation for 15 minutes. A 20 ul aliquot was taken for input, and the remaining cytoplasmic lysate was incubated with 240 ul Dynabeads MyOne Streptavidin C1 beads (Thermo Scientific, 65801D) for 45 minutes with gentle rotation. Beads were collected using a magnetic stand and gently resuspended in 1 ml modified RBB wash buffer (20 mM HEPES pH 7.5, 1. 5mM MgCl_2_, 0.5 M NaCl, 5% glycerol, 0.01% NP-40 and fresh 1 mM TCEP and protease inhibitor cocktail) before transferring to a 5 ml LoBind EP tube. Beads were washed twice with 4 ml modified RBB wash buffer for 15 minutes each with rotation, using a new 5 ml LoBind EP tube for each wash.

Pull-down beads were resuspended in 1 ml modified RBB wash buffer containing 2 ul TurboNuclease (Accelagen, N0103M) and transferred to 1.5 ml LoBind EP tube for overnight incubation at 4°C. The next morning, beads were washed twice with 1 ml modified RBB wash buffer by gentle pipetting and resuspended in 240 μl of modified RBB wash buffer for TRIM65 pull-down in the same day.

### TRIM65 pull-down

All the experiments were performed in RNase-free environment at 4°C or on ice unless otherwise indicated. Three 15 cm dishes of RPE-1 cells were used for each sample. For each dish, cells were first washed twice with 7 ml PBS and once with 4 ml hypotonic buffer (10 mM Tris pH 7.5, 10 mM KCl, 1.5 mM MgCl_2_) containing protease inhibitor cocktail (Roche, 11873580001). Cells were scraped in the residual 0.7-1 ml hypotonic buffer, transferred to a 1.7 ml Eppendorf tube, and gently resuspended with a 1 ml pipette, followed by 15 minutes of incubation. A 1 ml syringe with a 25G needle was used to gently pass the hypotonic buffer-resuspended cells through 7 times to lyse cells, avoiding bubbles. Cell lysates were centrifuged at 1,000 g for 5 minutes. The supernatant was transferred to a new tube, and NP-40 was added to a final concentration of 0.01%. The sample was then sequentially centrifuged at 5,000 g and 18,000 g for 5 minutes each. The supernatant containing cytoplasmic lysate from three dishes was carefully transferred and pooled into a 5 ml LoBind Eppendorf tube (Eppendorf, 0030108302) to achieve ∼1.5 ml, then mixed with an equal volume (1.5 ml) of RBB buffer (20 mM HEPES pH 7.5, 1. 5mM MgCl_2_, 150 mM NaCl) with 0.01% NP-40 containing protease inhibitor cocktail. The mixture was precleared with TrueBlot Protein G magnetic beads (Rockland, PG00-18-2) by gentle rotation for 15 minutes. A 100 ul aliquot was taken for input, and the remaining cytoplasmic lysate was incubated with 30 ul TRIM65 conjugated on magnetic beads for 45 minutes with gentle rotation. Beads were collected using a magnetic stand and gently resuspended in 1 ml RBB buffer with 0.01% NP-40 before transferring to a 1.5 ml LoBind EP tube (Eppendorf, 0030108442). Beads were washed twice with 1 ml RBB buffer with 0.01% NP-40 for 15 minutes each with rotation, using a new 1.5 ml LoBind EP tube for each wash.

Pull-down beads were resuspended in 1 ml low-salt RBB buffer (20 mM HEPES pH 7.5, 1. 5 mM MgCl_2_, 50 mM NaCl) with 0.01% NP-40. RNase A (Qiagen, 19101) was added to 1 ml resuspended pull-down beads and 100 μl input cytoplasmic lysate to a final concentration of 1 mg/ml, followed by incubation at 4°C for 15 minutes. RNase A-digested beads were washed twice with 1 ml RBB buffer with 0.01% NP-40 by gentle pipetting and then resuspended in 300 ul RBB buffer with 0.01% NP-40. Aliquots of 20 μl and 280 μl were taken for pull-down western blot analysis and RNA-seq, respectively.

For western blot, 20 μl cytoplasmic lysate and 20 μl Beads were boiled with 7 μl 4x NuPAGE LDS Sample Buffer (Thermo Scientific, NP0007) containing 10% 2-mercaptoethanol at 94°C for 5 minutes. For RNA-seq, an equal volume of 2x DNA/RNA shield (Zymo) was added to both the input lysate (50 μl) and the beads suspension (280 μl), followed by pipetting to mix. Proteinase K (Thermo Scientific, 25530049) was added to a final concentration of 1 mg/ml and incubated at room temperature for 30 minutes. An equal volume of DNA/RNA lysis buffer was then added to the input lysate-DNA/RNA Shield mixture (100 μl) and the beads suspension-DNA/RNA Shield mixture (560 μl), mixed by pipetting, and incubated at room temperature for 10 minutes. Beads were collected using a magnetic stand, and the supernatant was saved for RNA purification.

### RNA purification for TRIM65 pull-down

RNA for RNA-seq library preparation was purified using Quick DNA/RNA Kit (Zymo, D7005) according to manufacturer’s manual. Briefly, input RNA and eluted TRIM65 pull-down RNA that had been mixed with DNA/RNA shield and DNA/RNA lysis buffer, were combined with an equal volume of ethanol and passed through a Zymo-Spin IC column by centrifuge. The column was washed with DNA/RNA wash buffer, followed by on column DNase I digestion for 15 minutes at room temperature. After digestion, column was washed once with DNA/RNA prep buffer and twice with DNA/RNA wash buffer before elution with 20 ul of 65°C nuclease free water. RNA concentration was initially measured by a NanoDrop spectrophotometer and diluted accordingly for validation on an Agilent High Sensitivity RNA ScreenTape using a TapeStation 4150 (Agilent Technologies).

### RNA-seq library preparation

RNA-seq libraries were prepared using the SMARTer® Stranded Total RNA-Seq Kit v3-Pico Input Mammalian (Takara Bio, 634486) and SMARTer® RNA Unique Dual Index Kit-96U Set A (Takara Bio, 634452) according to manufacturer’s manual. In brief, 10 ng of input RNA and all pull-down RNA (since the total RNA amount was less than 10 ng) were used for library preparation. For non-RNase A treated samples, fragmentation was performed at 94°C for 4 minutes, and for RNase A digested samples, fragmentation was omitted before first strand cDNA synthesis. Illumina adapters with barcodes were added through PCR amplification: 5 cycles for non-RNase A treated samples and 10 cycles for RNase A treated samples, followed by ribosomal cDNA depletion. The final cDNA libraries were amplified by 13 PCR cycles for non-RNase A treated samples and 16 PCR cycles for RNase A treated samples, then purified using NucleoMag beads (Takara, 744970.5). cDNA library concentrations were initially measured by nanodrop and diluted accordingly for validation on an Agilent high sensitivity D1000 ScreenTape using a TapeStation 4150 (Agilent Technologies).

cDNA libraries were normalized to a concentration of 6 nM prior to pooling and pooled in an equimolar amount. The library pools were clustered and sequenced on a NovaSeq X platform with 150-base paired-reads, using sequencing services provided by Innomics.

### *De Novo* identification of A-to-I RNA editing sites

We performed *de novo* identification of A-to-I RNA editing sites in DDX3X depleted (DDX3X-mIAA7) and control samples separately, using the same pipeline and filtering criteria as described before^77,78^. We used UnifiedGenotyper tool from the Genome Analysis Toolkit (GATK)^79^ to call variants from RNA-seq reads mapped by STAR^80^. To maximize the accuracy of our site discovery, we applied previously established parameters and filters to these variant candidates. Specifically, variants located in human *Alu* regions required the support of at least one mismatched read, while those in non-repetitive or non-*Alu* repetitive regions required a minimum of three mismatched reads. To distinguish true RNA editing events from genomic polymorphisms, we filtered out known single nucleotide polymorphisms (SNPs) using data from the 1000 Genomes Project, the University of Washington Exome Sequencing Project, and dbSNP (version 135, explicitly excluding ‘cDNA’ molecular types). Furthermore, we conducted a hyper-editing analysis to rescue highly edited sites from the pool of unmapped reads using SPRINT^81^. Finally, the newly identified editing sites from each cell line were merged with known annotated sites from the RADAR database^82^.

### Identification of A-to-I RNA editing clusters

To identify the dsRNAs that are potential MDA5 ligands without relying on any prior functional annotation, we focused on hyper-edited regions as highly immunogenic MDA5 ligand candidates. The location information of the editing sites was clustered using a density-based clustering algorithm (DBSCAN) to identify editing clusters, which represented proxies for dsRNA regions as described before^24^. We used these two parameters to define a cluster: the minimum number of points (editing sites) required to form a cluster (minPts ≥ 5) and the farthest distance between two adjacent sites within the same cluster (eps ≤ 120).

### Analysis of RBP binding and pull-down data

TRIM65-pulldown peaks were called on crosslinked nucleotides with JAMM (version 1.0.7rev6, parameters: -d y -t paired -b 50 -w 1 -m normal)^83^ using two replicates of the respective pull-down as foreground and the two replicates of the corresponding input condition as background. The peaks called in the knockout of MDA5 were treated as non-specific background signals and subtracted from the peaks called in the presence of MDA5. CLIP-seq data for ADAR1^51^, DDX3X^50^, uvCLAP data for DHX9^20^ and J2 (dsRNA) pull-down data^20^ were analyzed with PEAKachu (version 0.2.0, parameters: –pairwise_replicates -m 0 -n manual)^84^, using at least two replicates of the respective pulldown as foreground and at least two replicates of the corresponding input as background. The peaks of all experiments were merged and used to calculate pairwise Pearson correlations based on the number of events falling on each peak region using deepTools (version 3.5.6)^85^.

To determine the enrichment of repetitive elements in pulldown samples compared to input controls, we employed a previously established mapping-free, graph-based sequence clustering method^20,86^. Briefly, we sampled 1,000,000 reads from the MDA5, ADAR, DDX3, DHX9, and J2 binding profiling datasets. These reads were processed through a clustering pipeline^87^ that conducts all-to-all pairwise comparisons to construct a sequence graph. In this graph, reads serve as vertices (nodes), and edge weights reflect the sequence similarity scores between them. The graph was subsequently partitioned into distinct clusters representing connected communities. Next, we utilized RepeatMasker to annotate the reads within each cluster for the presence of various repeat families. Finally, the largest clusters (those with the highest read counts) for each sample were visualized to evaluate the composition of major Alu families and other repetitive elements.

### Characterization of dsRNA features

To characterize the features of dsRNAs, we first predicted minimum free energy (MFE) secondary structures using the RNAfold tool from the ViennaRNA package (default parameters)^88^. For these predictions, the analyzed sequences comprised the closest inverted pairs along with their intervening non-repeat regions. We then computed the partition function to determine both the ensemble diversity and the frequency of the MFE structure within the thermodynamic ensemble. Additional features of dsRNAs were also obtained from dsRNAscan^89^ for comparisons. Based on the predicted and annotated dsRNA features, we quantified and compared structural properties, including the fraction of paired bases and the total number of mismatches and bulges.

### Differential RNA editing analysis

For cancer data, patients from the TCGA cohort with DDX3X somatic mutations, as identified and specified by Yanas et al.^36^ showing potentially association with RNA unwinding, were selected for analysis only if both tumor and matched solid normal samples were available. RNA editing data was obtained from the GPEdit database^54^. RNA-seq data of mouse medulloblastoma with DDX3X depletion was obtained from GEO: GSE147069^55^.

For differential RNA editing analysis, only editing sites with a minimum coverage of 20X were included for comparison. For each editing site meeting the above threshold in both conditions in comparison, Fisher’s exact test was applied to the counts of G and A reads to calculate the P value to test if editing levels are equal. Multi-test p-values were adjusted using the Benjamini–Hochberg procedure.

### Statistical Analysis

Unless other specified, all statistical analyses were performed using GraphPad Prism. Quantitative data are presented as mean ± SEM, and statistical significance was assessed using two-tailed unpaired Student’s t test.

---

**Extended Data Fig. 1 Generation and validation of DDX3X degron cells and analysis of innate immune signaling following ATR inhibition and irradiation. a**, Cells were infected with lentiviral guide RNAs targeting indicated genes. 4 days post infection, cells were treated with 2.5 µM ATRi for 1 hour before 10 Gy irradiation (IR) or no irradiation (NIR). Cells were then cultured for an additional 3 days in the presence of 2.5 µM ATRi before western blot analysis. **b**, Schematic of the workflow used to generate DDX3X degron bulk polyclonal cells, including CRISPR–Cas9 targeting of the DDX3X locus, donor plasmid insertion, GFP-based enrichment, and recovery of homozygous degron cells. **c**-**d**, Validation of degron tag insertion and auxin-induced degradation of DDX3X. **c**, Parental and DDX3X degron (DDX3X-mIAA7) cells were subjected to western blot analysis to validate degron tag insertion. **d**, Time-course of DDX3X degradation following treatment with 1 mM auxin for the indicated durations followed by western blot analysis. **e**, Cells were treated with 2.5 µM ATRi for 1 hour, followed by 10 Gy irradiation (IR) or no irradiation (NIR), and cultured for 3 days in the presence of 2.5 µM ATRi prior to western blot analysis. **f**, Cells were treated with 2 µg/ml DOX for 24 hours, then incubated for 3 hours with 2 µg/ml DOX, 1 mM auxin and 2.5 µM ATRi before 10 Gy irradiation (IR) or no irradiation (NIR). Cells were cultured for an additional 2 days in the same drug combination prior to western blot analysis.

**Extended Data Fig. 2 DDX3X depletion elevates cytosolic dsRNA and synergizes with DNA damage or ADAR1 deficiency to promote robust ISG induction. a**, Cells were fixed with paraformaldehyde (PFA) and permeabilized with saponin prior to dsRNA immunostaining using the 9D5 and J2 antibodies. **b**, **e**-**h**, Cells were treated with 1mM auxin in the presence of 2.5 µM ATRi for 3 hours, followed by 10 Gy irradiation (IR) or no irradiation (NIR). Cells were then cultured for another 3 days in the presence of 1 mM auxin and 2.5 µM ATRi before immunostaining analysis (**b**) or flow cytometry analysis (**e**-**h**). **c**-**d**, Cells were infected with lentiviral guide RNAs targeting indicated genes. 4 days post infection, cells were treated with 1 mM auxin for 2 days before flow cytometry analysis (**c**) or western blot analysis (**d**). Panel (**c**) corresponds to the quantification of **Fig.1g**.

**Extended Data Fig.3 Genomic location and repeat composition of edited dsRNAs. a**, Transcript feature annotations of edited dsRNAs detected in cytosolic RNA from parental and DDX3X depleted (DDX3X-mIAA7) cells, showing the proportion of edited dsRNAs mapping to 5’ untranslated regions (5’UTRs), 3’ untranslated regions (3’UTRs), coding sequences (CDS), introns, and promoters. **b**, Repeat sequence composition of edited dsRNAs in control cells, classified by repeat classes. **c**, Repeat sequence composition of edited dsRNAs in DDX3X-depleted (DDX3X-mIAA7) cells relative to control (DDX3X-mIAA7 minus control), classified by repeat classes.

**Extended Data Fig.4 Characterization of TRIM65 pull-down specificity and associated dsRNAs. a**, Cells were transfected with a plasmid expressing SBP-tagged TRIM65 and cultured for 3 days prior to TRIM65 purification. Silver staining was performed to validate TRIM65 expression and purity. **b**, **d**, Cells were infected with lentiviral guide RNA targeting indicated genes. 4 days post infection, cells were treated with 1 mM auxin for 2 days prior to TRIM65 pull-down assay, followed by western blot analysis (**b**) and stranded RNA-seq prior to analysis of the relative contribution of repeat classes in pull-down background dsRNAs recovered from DDX3X-mIAA7 + ADAR1 KO + MDA5 KO cells (**d**). **c**, Cells were treated with 1mM auxin in the presence of 2.5 µM ATRi for 3 hours, followed by 10 Gy irradiation (IR) or no irradiation (NIR). Cells were then cultured for another 3 days in 1 mM auxin and 2.5 µM ATRi before western blot analysis.

**Supplementary Table 1.**
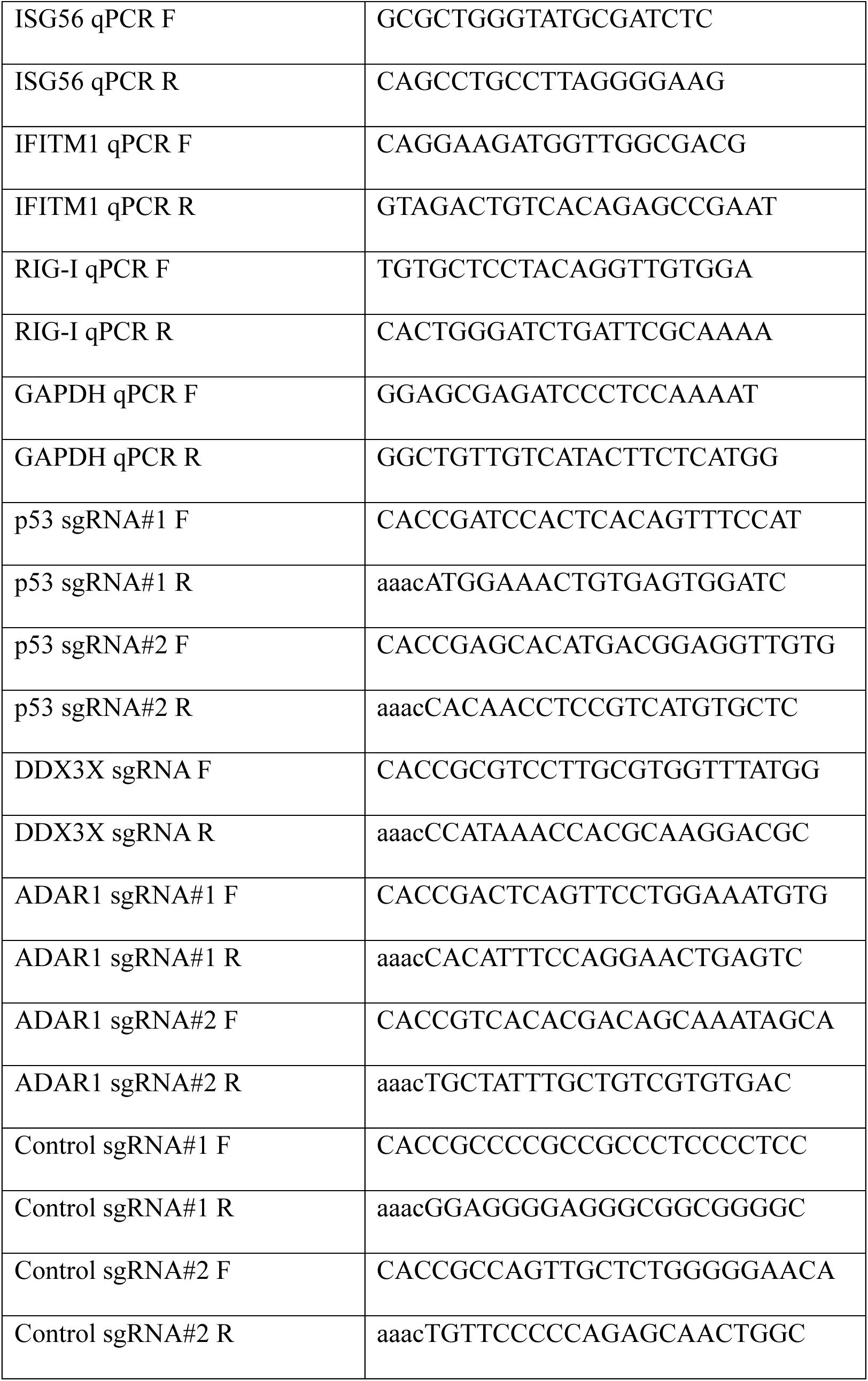
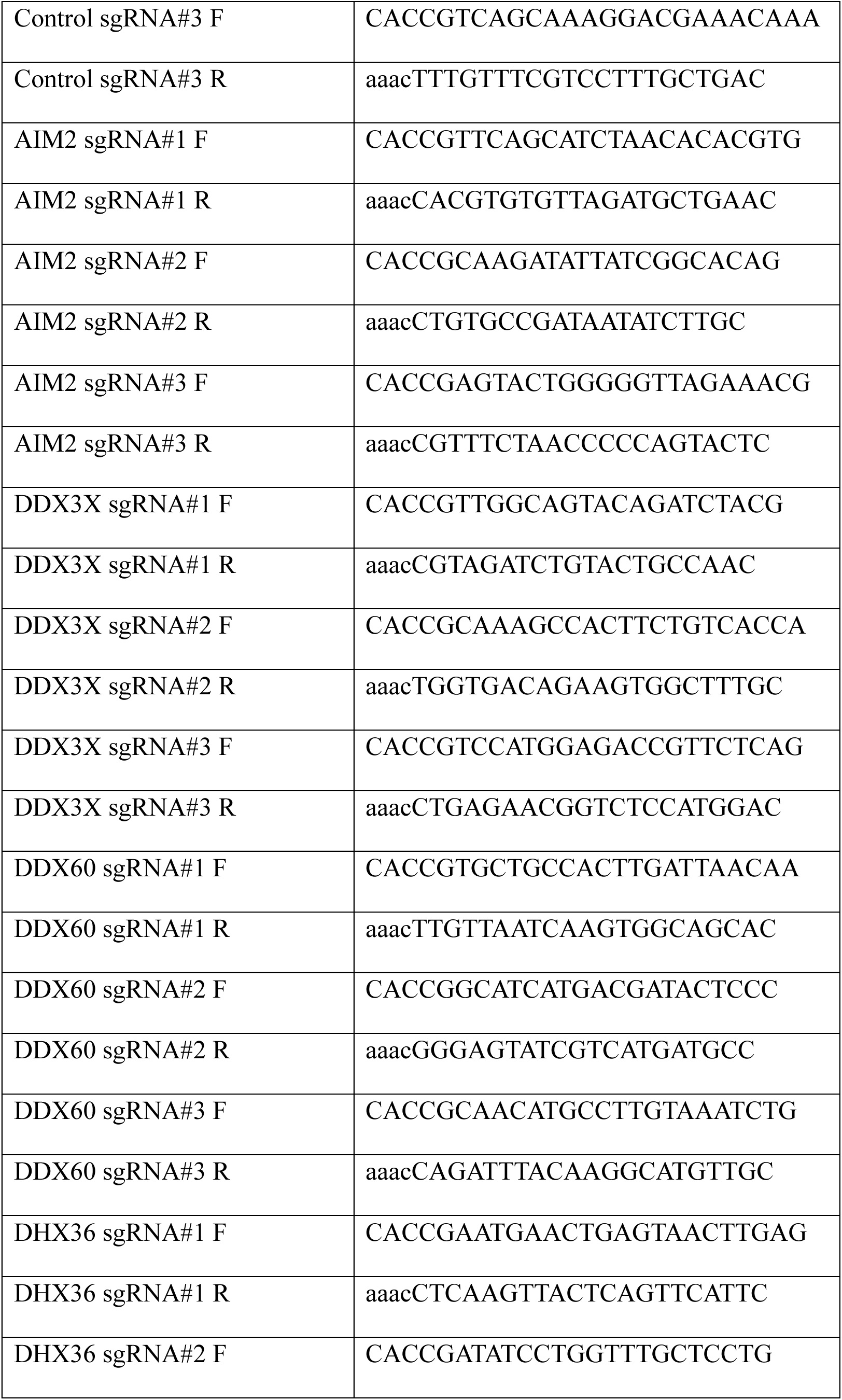
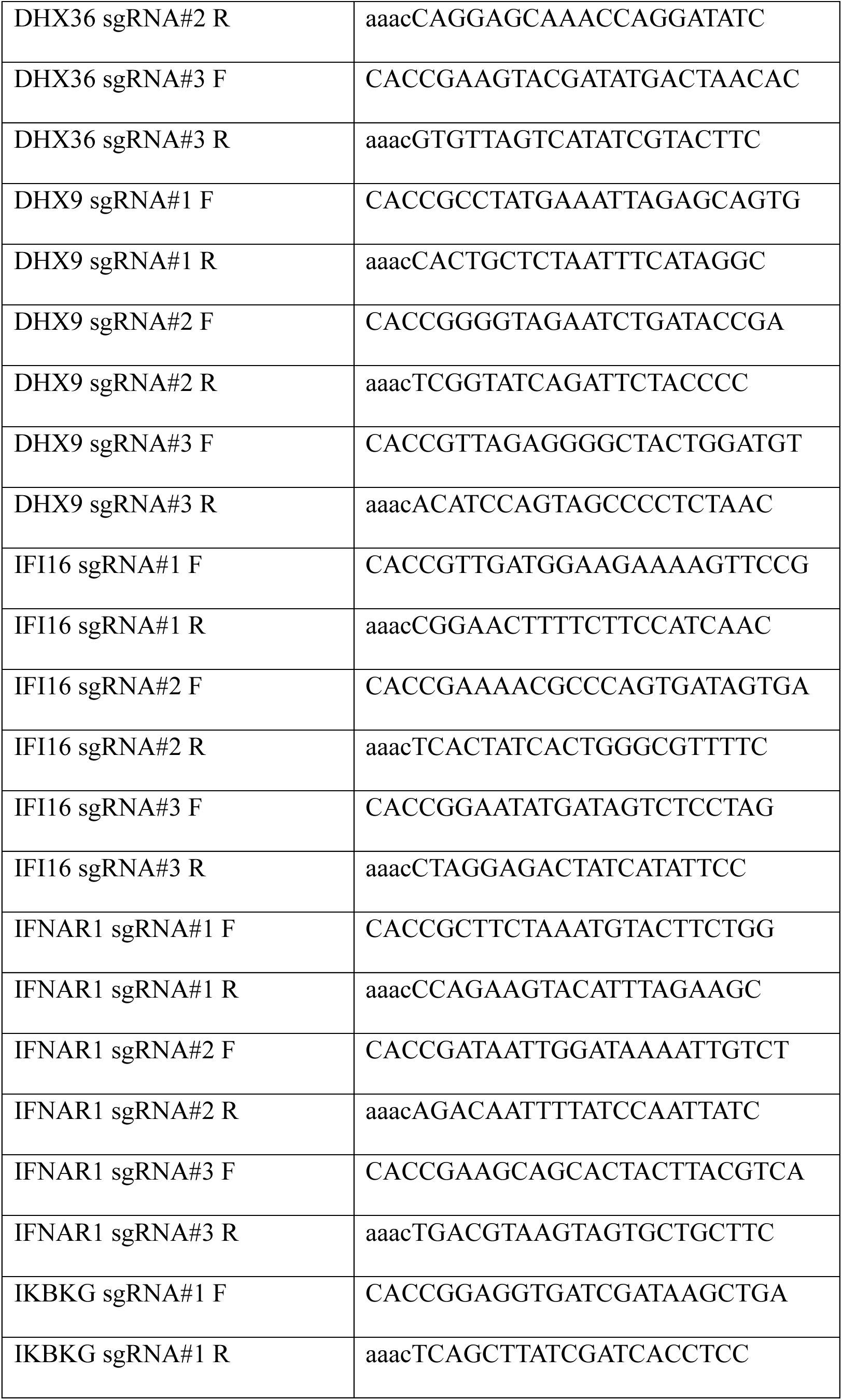
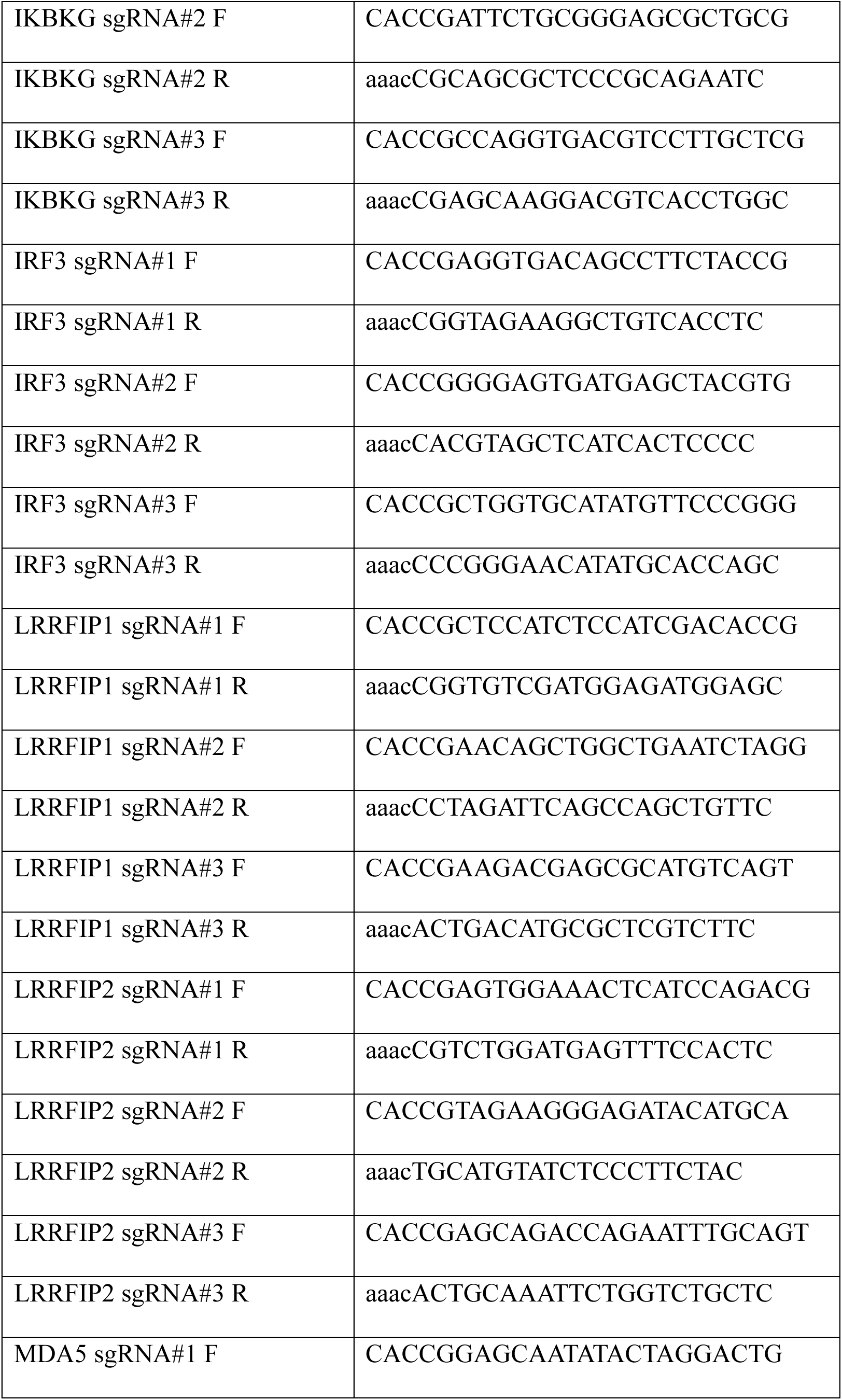
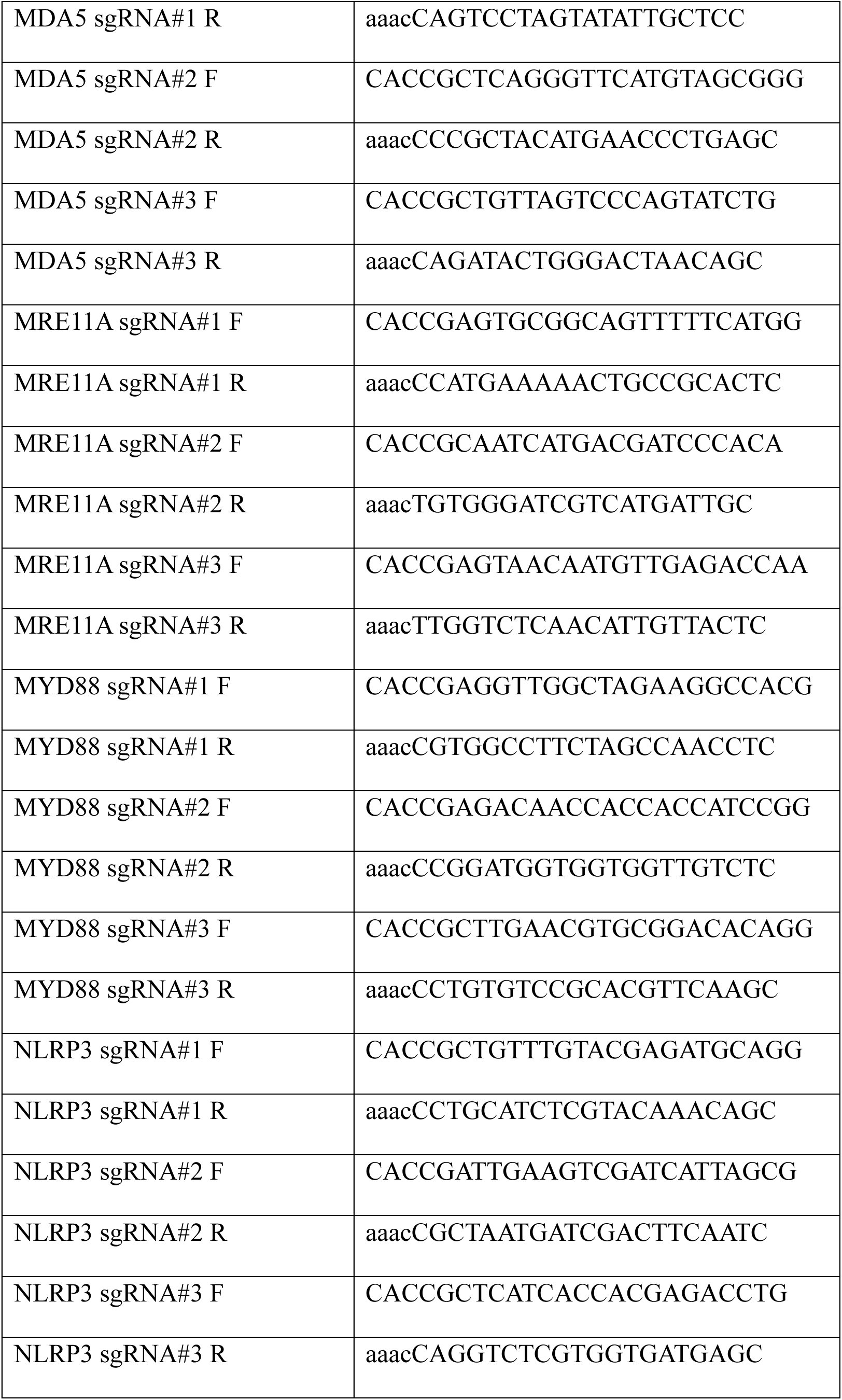
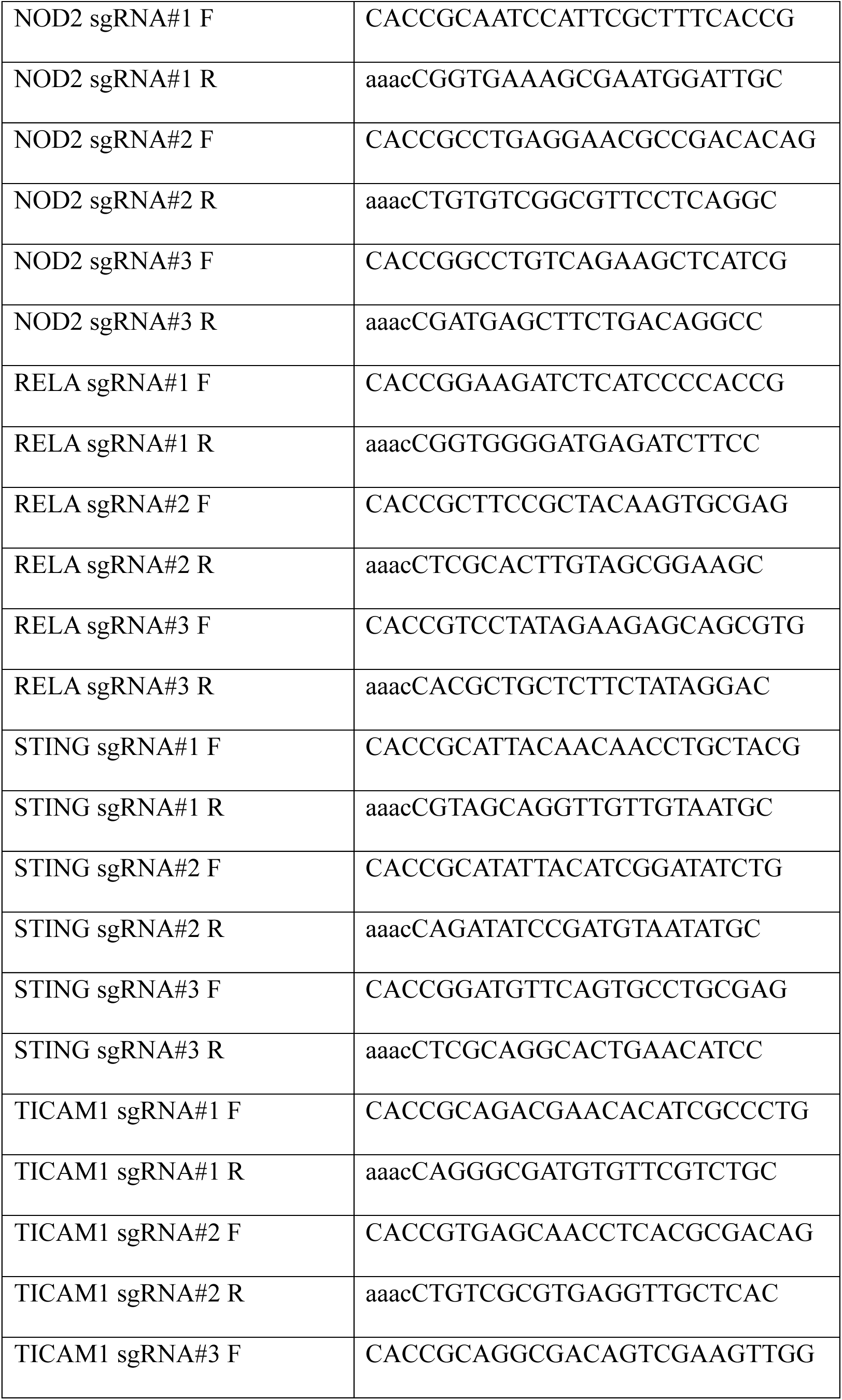
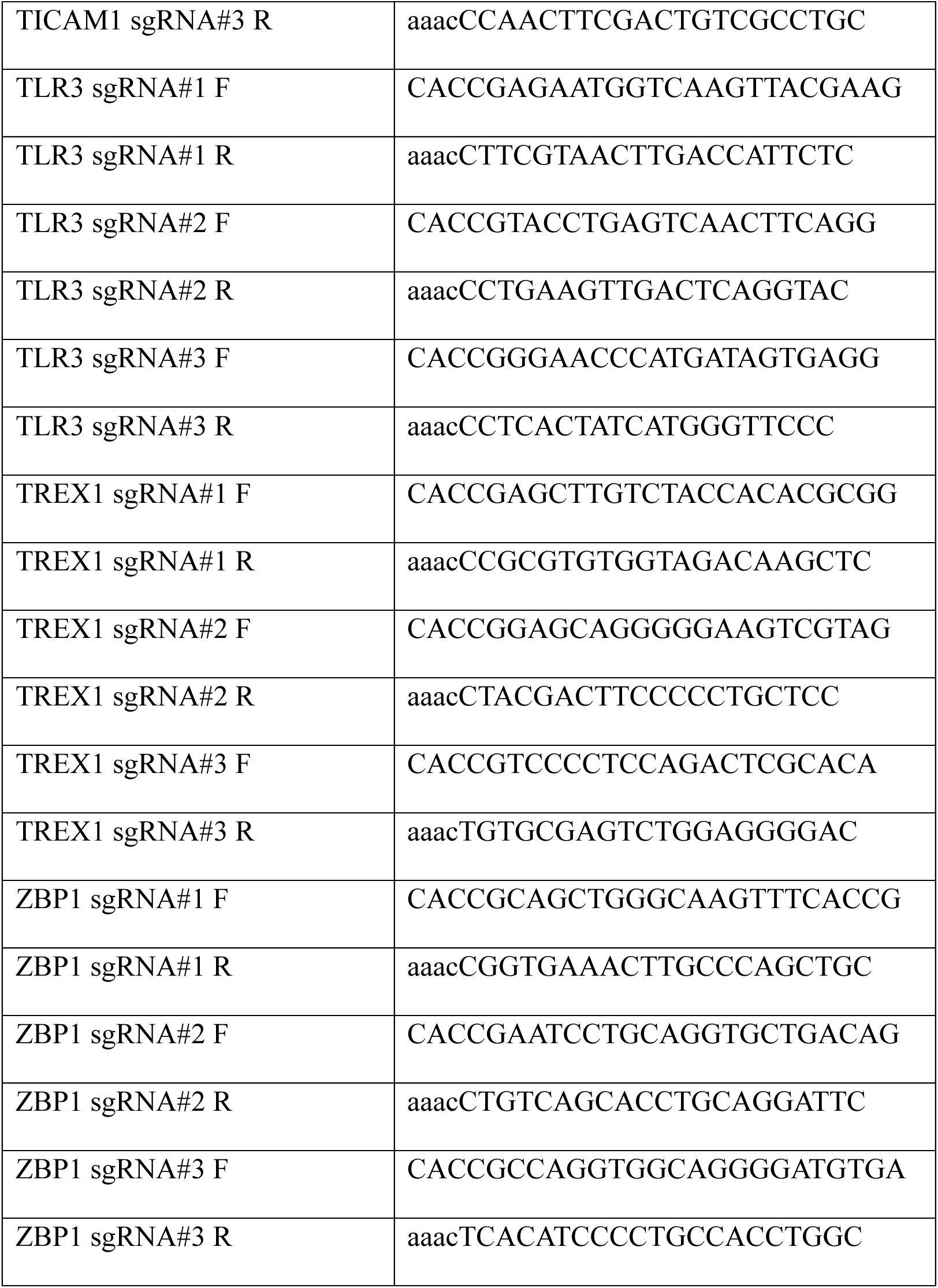
Oligonucleotides.

## Acknowledgments

We thank members of the Greenberg, Li, Hur, and Liu labs for critical discussion. This work was supported by NIH P01 CA265794 and R01 CA174904 from the NCI (R.A.G.), U01DK143477 from NIDDK and R35GM162298 from NIGMS (Q.L.), the MARK Center for Radiation Oncology (R.A.G.), and the Penn Center for Genome Integrity (R.A.G., Q.L., and K.F.L). J.C. is the recipient of an AFCRI RISE Fellowship (UPENN). (Research reported in this publication was supported by the National Cancer Institute of the National Institutes of Health. The content is solely the responsibility of the authors and does not necessarily represent the official views of the National Institutes of Health.)

## Competing interests

The authors declare no competing interests.

**Correspondence and requests for materials** should be directed to and will be fulfilled by Qin Li and Roger Greenberg.

