## Supplemental Figures for "Identification of highly immunogenic endogenous dsRNAs from cellular MDA5 filaments"

**Extended Data Fig.1 Generation and validation of DDX3X degron cells and analysis of innate immune signaling following ATR inhibition and irradiation**

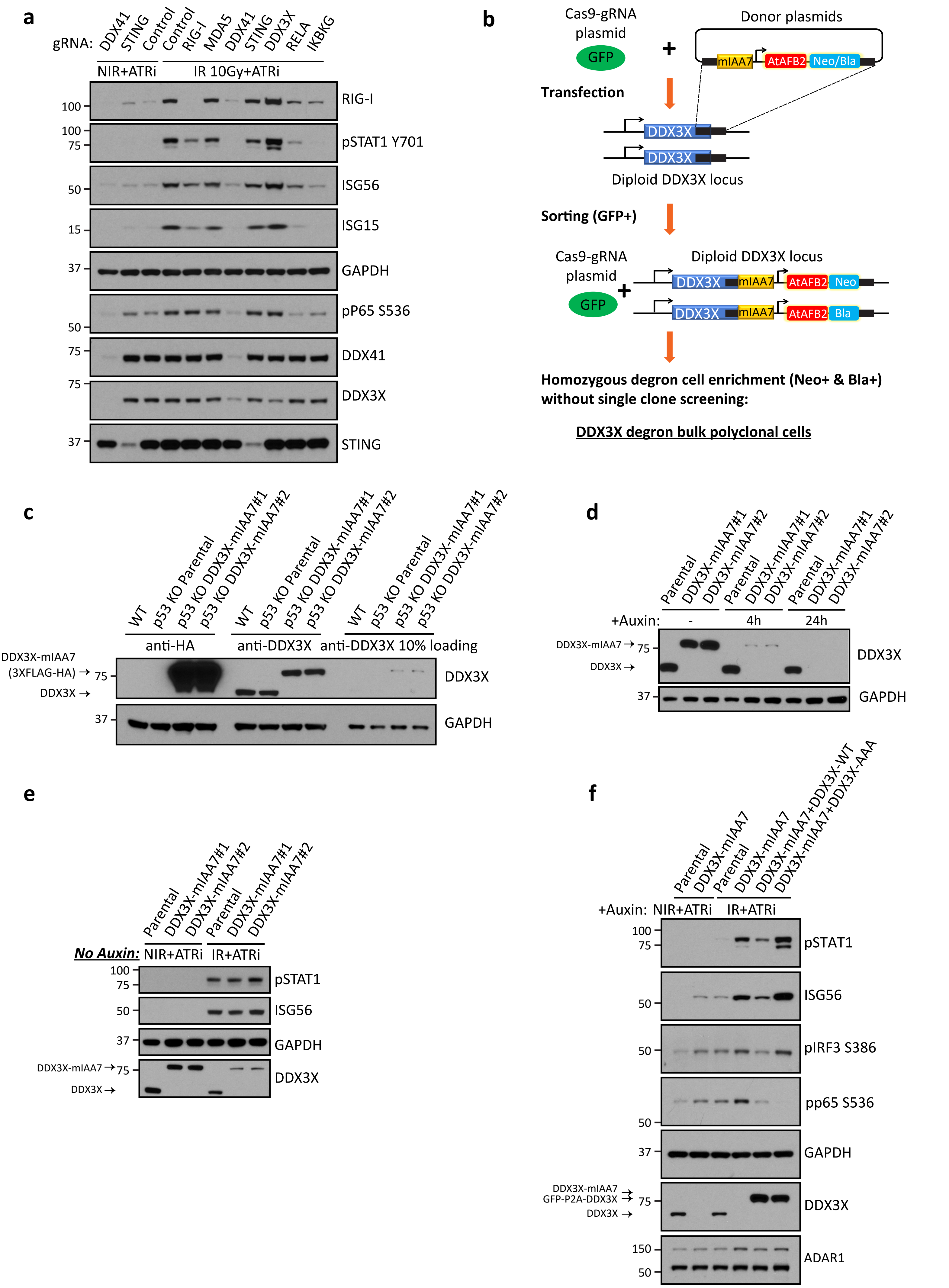

#### Extended Data Fig.2 DDX3X depletion elevates cytosolic dsRNA and synergizes with DNA damage or ADAR1 deficiency to promote robust ISG induction

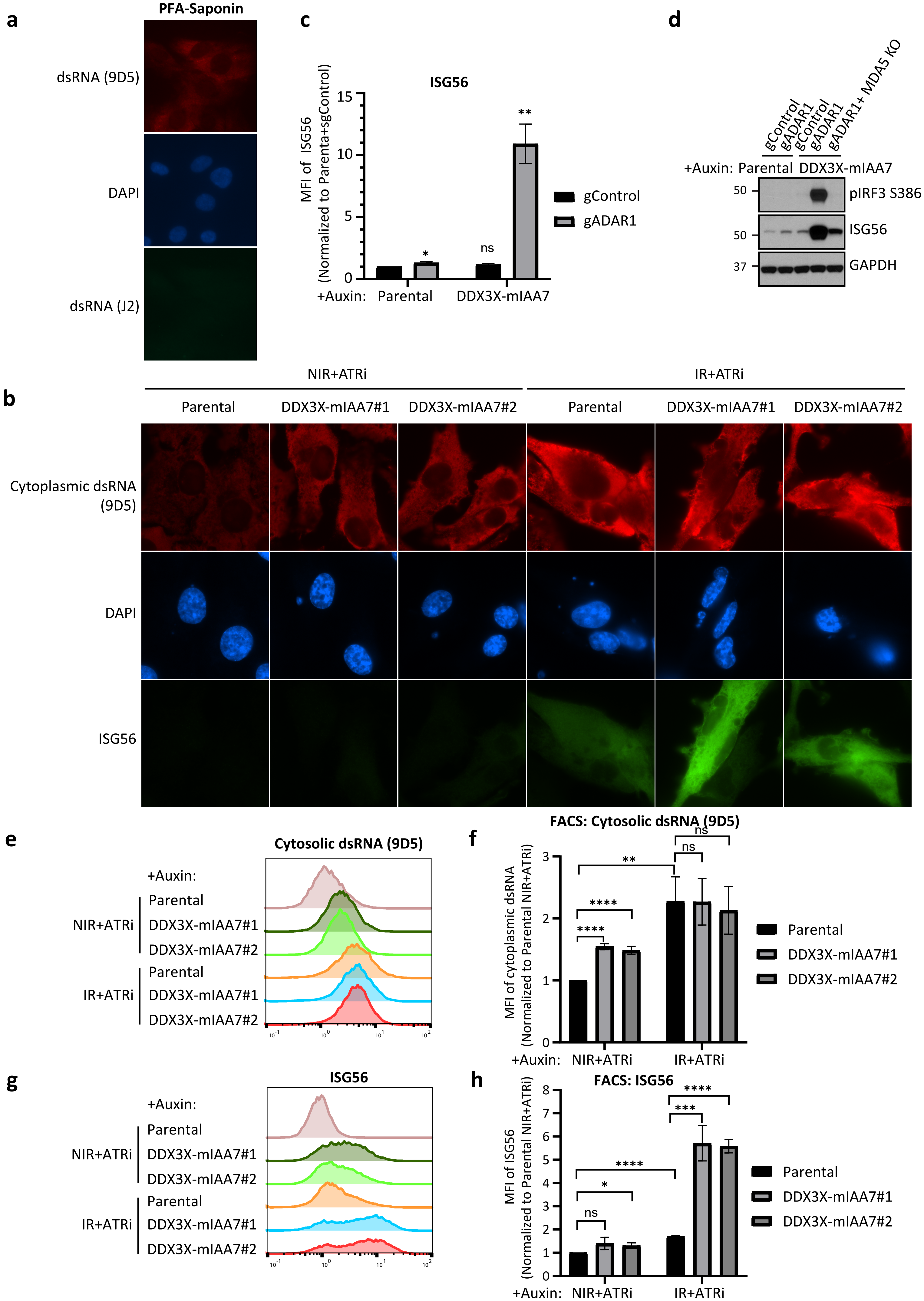

### Extended Data Fig.3 Genomic location and repeat composition of edited dsRNAs

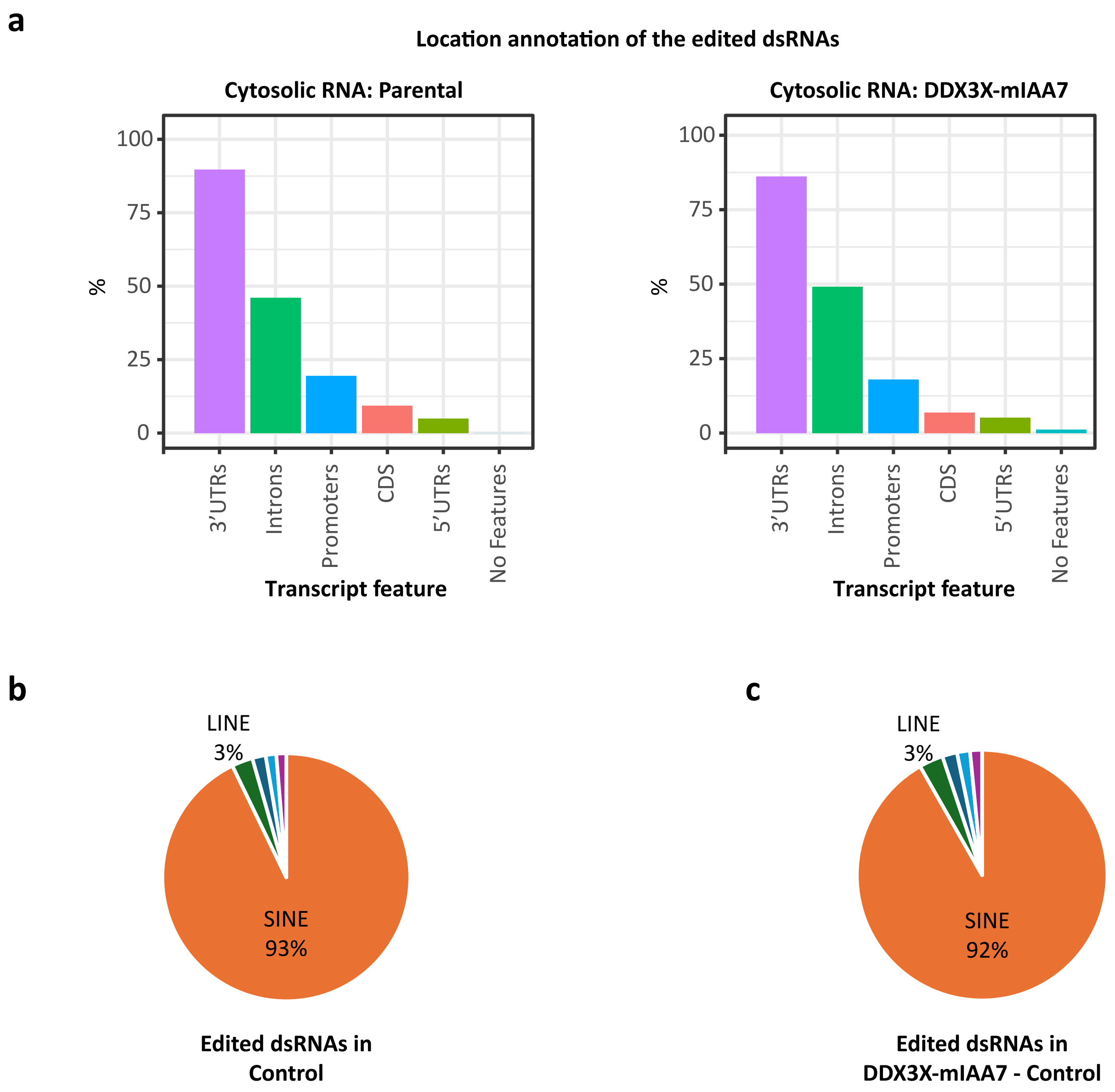

#### Extended Data Fig.4 Characterization of TRIM65 pull-down specificity and associated dsRNAs

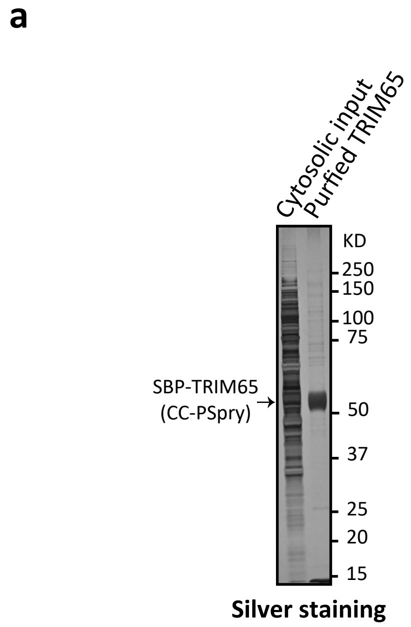


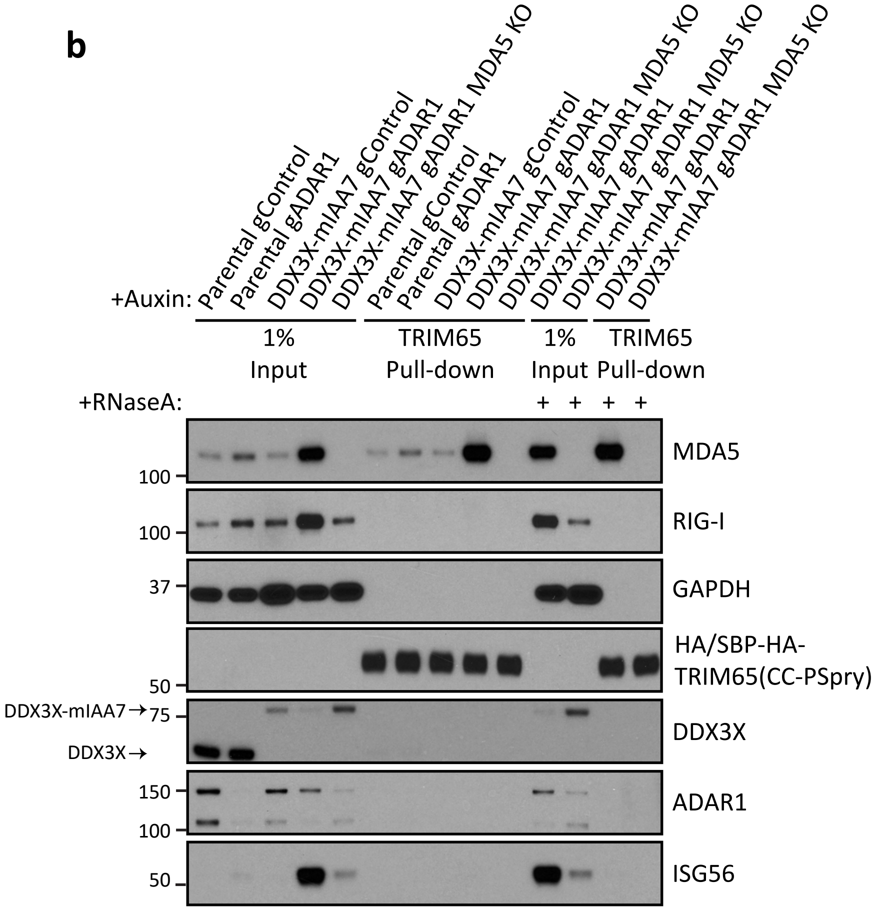


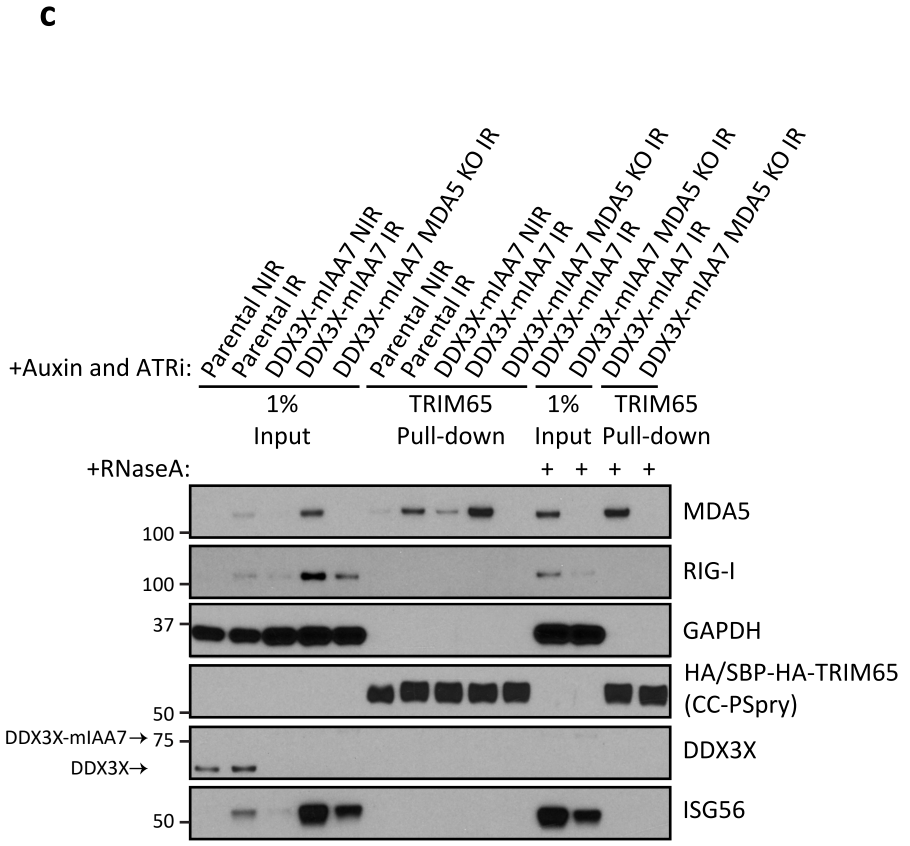


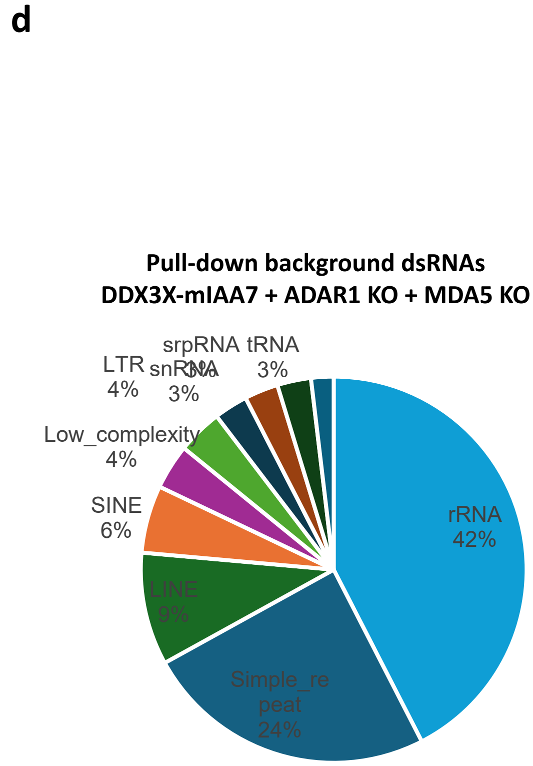
